# CRISPR-activation reveals key resistance genes and vulnerabilities of copy number variants in *Candida albicans*

**DOI:** 10.64898/2026.08.07.743545

**Authors:** Pétra L Vande Zande, Nicholas C Gervais, Erika Schell, Piper Zajac, Katura Metzner, Rebecca S Shapiro, Anna Selmecki

## Abstract

Changes in gene copy number are an extremely important source of variation and are frequently observed during the acquisition of drug resistance. The opportunistic human fungal pathogen *Candida albicans* frequently adapts to antifungal drugs via large copy number variations (CNVs) that amplify hundreds of genes simultaneously. Despite the recurrent amplification of CNVs across diverse clinical isolates, the genes that contribute to drug resistance are not known. Additionally, by amplifying many genes, CNVs might result in cross-adaptation or fitness trade-offs to multiple environments, which has major implications for how CNVs are expected to contribute to adaptation in complex environments like a mammalian host. We use CRISPR-activation to systematically assay the fitness effects of individually overexpressing ∼800 genes in four genetically diverse isolates across eight physiologically relevant environments. We identify 198 genes with significant fitness effects in at least one environment in one or more genetic backgrounds. We identify novel genes with positive fitness effects in two different classes of antifungal drug and observe frequent gene-by-environment interactions for the fitness effects of gene overexpression. Additive fitness effects of individual gene overexpression are a significant predictor of the fitness of multiple isolates with CNVs and can explain fitness trade-offs observed between classes of antifungal drug for the CNV isolates. These findings identify genes that increase fitness in drug and those that create vulnerabilities in CNV isolates and can help inform treatment of isolates adapting to antifungal drug via CNVs.

## INTRODUCTION

Copy number variants (CNVs) are important sources of phenotypic variation across kingdoms and frequently contribute to microbial adaptation to drugs and environmental stressors.^1–6^ Large CNVs can affect hundreds of genes and, on average, expression levels increase in proportion to copy number.^7–10^ Delineating the effects of gene amplification events that can impact expression of hundreds of genes simultaneously is a critical challenge in genetics. The fitness effects of CNVs are often environment- and CNV-specific and likely driven by the changes in expression of the genes encoded within the CNV.^11–13^ Because CNVs impact expression of many genes, one CNV might result in cross-adaptation or fitness trade-offs across different environments. These potential pleiotropic effects of CNVs have major implications for how they are expected to contribute to adaptive evolution in complex and changing environments such as the mammalian host.^14–17^ Nevertheless, how each gene within a CNV contributes to the fitness effect of a CNV in one environment, much less multiple environments, is not well understood.

Identifying the genes that contribute to the fitness effects of CNVs in any environment is challenging. Studies typically rely on existing functional annotations to identify candidate genes within large CNV regions that might contribute to a phenotype,^18–22^ leaving the contributions of uncharacterized genes unexplored. In addition, many adaptive CNVs increase gene copy number, resulting in overexpression of encoded genes, but most annotations of gene function are based on loss of function phenotypes. Furthermore, in the model organism *Saccharomyces cerevisiae,* the fitness effects of gene overexpression are dramatically impacted by genetic background^23^ and often exhibit gene-by-environment interactions.^19^ For these reasons, identification of genes within CNVs that contribute to fitness is a major challenge, particularly across genetically diverse backgrounds of organisms with many uncharacterized genes.

The opportunistic fungal pathogen *Candida albicans* has an incredibly plastic genome. Clinical isolates frequently exhibit aneuploidy and CNVs of different lengths and amplitudes affecting hundreds of genes.^20,21,24–31^ DNA copy amplifications are associated with azole antifungal drug tolerance and resistance, underscoring the need to understand gene amplification during adaptation.^5,21,30–34^ In rare cases, individual genes within these CNVs have been identified as drivers of drug resistance. For example, amplification of the left arm of chromosome 5 in an isochromosome structure (i(5L)) is both necessary and sufficient to cause resistance due to an increase in copy number of the drug target, *ERG11*, and a transcriptional regulator of drug efflux pump genes, *TAC1.*^35^ Similarly, amplification of *NCP1* in a CNV on chromosome 4 leads to decreased azole sensitivitiy^36^ and strains adapted to hydroxyurea via a chromosome 2 trisomy were cross-adapted to both hydroxyurea and caspofungin.^18^ Notably, different genes within chromosome 2 lead to decreases in susceptibility to each drug,^18^ demonstrating that multiple genes affected by large CNVs can result in cross-adaptation to multiple drugs. In each of these cases, the genes contributing to the phenotypes were identified from previous annotation, and whether additional genes contribute to the phenotypes observed is unknown.

*In vitro* adaptation to azole antifungal drugs leads to the rapid acquisition of complex CNVs in *C. albicans*^20,37,38^ similar to those observed in clinical isolates of *C. albicans* and other species.^6,25,26,28,29^ Complex CNVs are comprised of stair-step amplifications, consisting of a high copy central region with adjacent lower copy regions, each of which are flanked by long inverted repeats (Extended Data Fig. 1, Supplementary Table S1).^20,38^ The CNVs are associated with increased minimum inhibitory concentration (MIC_50_) to fluconazole and arose *de novo* in diverse genetic backgrounds. Multiple CNVs affected regions on chromosomes 1, 3, and 4 in *C. albicans*. The identity of the genes contributing to decreases in susceptibility and how they influence fitness in other environmental conditions or antifungal drugs is unknown.

Here we use CRISPR-activation to systematically measure the fitness effects of individual gene overexpression and relate them to the fitness effects of CNVs. We identify genes amplified in CNVs that have effects on fitness in fluconazole and six other physiologically relevant environments across four genetically diverse isolates. We find that gene-by-environment interactions are prevalent, but that genes with larger effect sizes are also more consistent across genetic backgrounds. Most fitness effects of individual genes are environment specific. Therefore, different genes within CNV regions contribute most strongly to fitness in each environment. We found a simple additive model of individual gene overexpression fitness to be a significant predictor of the fitness effects of multiple strains with CNVs, including fitness trade-offs between antifungal drug classes. The additive model predicts that the fitness trade-off between drug classes depends on the precise boundaries of the CNVs, but that refining a CNV to resolve trade-offs may be unlikely due to the positions of long inverted repeats. These results support the idea that complex CNVs are likely to serve as transient sources of adaptation and will be either refined or eliminated over time in complex environments such as the human host.

## RESULTS

### CRISPR-activation induces target gene overexpression in genetically diverse isolates of C. albicans

To systematically assay the fitness effects of individual gene overexpression for genes within complex CNVs, we utilized a CRISPR-activation (CRISPRa) system^39,40^ in which a catalytically dead Cas9 fused to a transcriptional activator complexes with a single guide RNA (sgRNA) and results in transcriptional activation of the gene targeted by the sgRNA. A construct including all components of the system is integrated into the *C. albicans* genome at the neutral locus *NEUT5L* (Fig. 1A). We selected four genetically diverse isolates (AMS5192, L26, P75063, and P75016) of *C. albicans* that have extensive phenotypic characterization.^21,25,38,41–44^ AMS5192 is a prototrophic derivative of the reference SC5314-derived background Sn152. L26 differs from SC5314 by 5,414 homozygous SNPs and is within the same clade, while P75063 and P75016 are from a distant clade and differ from SC5314 by 40,138 and 31,516 homozygous SNPs, respectively.^26^

**Fig. 1:**
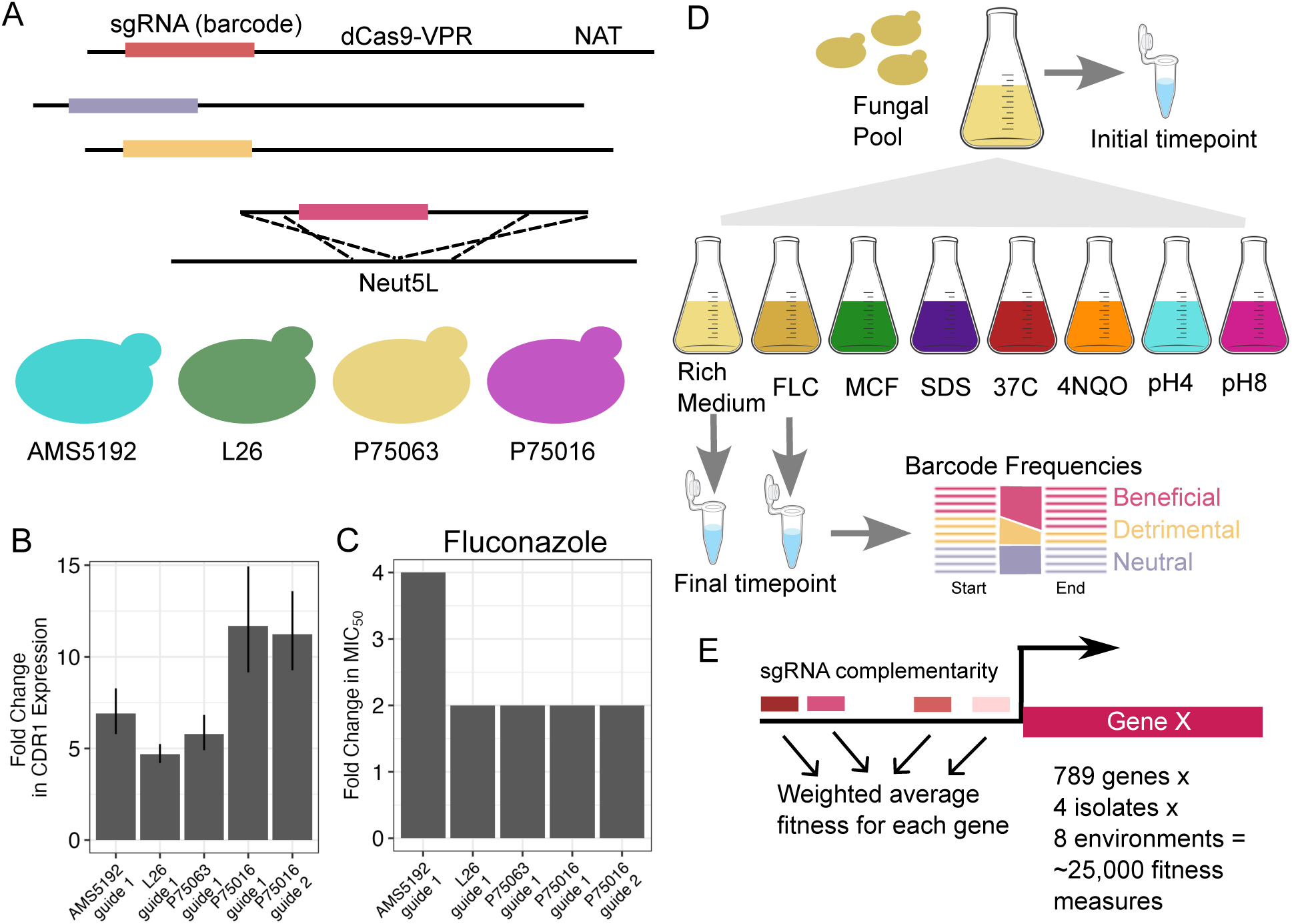
CRISPR-activation for measuring fitness in diverse genetic backgrounds and environments. (A) The CRISPRa system consists of dCas9-VPR and sgRNA along with a selectable Nourseothricin resistance (NAT) marker. The pooled constructs, each containing a unique sgRNA, are linearized and integrated into the genome at the *NEUT5L* locus. The pooled constructs were transformed into 4 different genetic backgrounds of *C. albicans*: AMS5192, L26, P75063, and P75016. (B) Change in *CDR1* expression relative to *ACT1* as measured by RT-qPCR in all four genetic backgrounds transformed with CRISPRa targeting *CDR1* with guide 1 or guide 2. (C) Fold-change in MIC_50_ in the same transformed strains as shown in (B) relative to the untransformed backgrounds. (D) Schematic of the pooled bulk competition assays to measure fitness of each barcoded genotype in each pool in 8 environmental conditions, including: 1) rich medium; 2) 1 μg/mL fluconazole (FLC); 3) 0.016 μg/mL micafungin (MCF); 4) 0.03% Sodium Dodecyl Sulfate (SDS); 5) 0.125 μg/mL 4-Nitroquinoline 1-oxide (4-NQO); 6) 37°C (all other conditions were performed at 30°C); 7) rich media adjusted to pH4 using HCl; and 8) rich media adjusted to pH8 using NaOH. In each genetic background for each of 3 replicates, the same initial inoculum was used to inoculate 8 different environmental conditions, and a portion of the inoculum was frozen down as the initial timepoint. After the competition, aliquots of each environmental condition were frozen down. Deep barcode sequencing of initial and final timepoints was then used to quantify changes in barcode frequency during competition. (E) To calculate fitness estimates for each gene overexpression, a weighted average of all guides targeting the same genes was calculated (see Methods and Extended Data Fig. 3).

We transformed each genetic background with a CRISPRa construct containing a sgRNA targeting the promoter of the efflux pump gene *CDR1*^39^ and performed broth microdilution assays in fluconazole. In all four genetic backgrounds, CRISPRa transformants had significantly higher expression of *CDR1* than the progenitor isolates by RT-qPCR and had 2- to 4-fold increased fluconazole MIC_50_ (Fig. 1B,C). These results demonstrate that the CRISPRa system induces overexpression of the targeted gene and results in a significant phenotypic change in all four genetic backgrounds.

### Pooled competition assays reveal fitness effects of overexpression across genetic backgrounds and environments

We next systematically assayed the fitness effects of individual gene overexpression for 843 genes located in regions of the genome that have been amplified via complex CNVs during evolution experiments (Extended Data Fig. 1, Chromosome 1:2117999 to 3133121, Chromosome 3L:621867 to 897198, Chromosome 3R:1047995 to 1457807, Chromosome 4:531821 to 898541). We generated sgRNA pools consisting of 2-10 sgRNAs targeting each gene and 60 non-targeting sgRNAs for a total of 5,885. We then generated pooled libraries, in which each cell contains one sgRNA, in each of the four genetic backgrounds. Deep sequencing of the sgRNA sequences integrated at *NEUT5L* showed high pool complexity in all four fungal pools, with only 8 guides absent from any pool (Extended Data Fig. 2).

To measure the fitness effects of individual gene overexpression we performed bulk competition assays of each CRISPRa pool under eight physiologically relevant environmental conditions performed in triplicate, including rich medium, fluconazole, micafungin, sodium dodecyl sulfate, 4-nitroquinoline 1-oxide, 37°C, pH4, and pH8 (Fig. 1D). We calculated the log_2_ fold change in frequency of each barcoded strain from the inoculum (time = 0) to 24 hours of competitive growth to calculate a fitness score for each barcoded strain. To calculate fitness estimates for each gene overexpression, we used a weighted average of all sgRNAs targeting the same gene (Fig. 1E, see Methods and Extended Data Fig. 3). After quality filtering we measured fitness for 789 genes in four genetic backgrounds and eight environmental conditions (Fig. 1E). We then calculated mean and standard deviation for each gene fitness score across the three replicates to identify genes with significant fitness effects when overexpressed.

### Fitness effects of overexpression in rich medium are small and on average negative

In all genetic backgrounds the mean fitness effects of gene overexpression in rich medium were negative (Fig. 2A). Genes with significant effects on fitness compared to non-targeting controls (moderated t-test with BH correction FDR <= 0.05) were almost entirely specific to each genetic background (Fig. 2B,C). One gene, *C3_05110W*, was significantly detrimental when overexpressed in all four genetic backgrounds (Fig. 2B,C). We generated independent transformants overexpressing *C3_05110W* in all four genetic backgrounds and performed growth curves in rich medium for each transformant and the progenitor strains. We found that growth rate in rich media was significantly reduced in strains overexpressing *C3_05110W* only in the AMS5192 genetic background (Fig. 2D), where it had the largest effect size in the pooled competition assay (Fig. 2B). Overall, these results demonstrate that gene overexpression in rich medium is on average deleterious but generally has small effect sizes that are not detectable in growth curve assays.

**Fig. 2:**
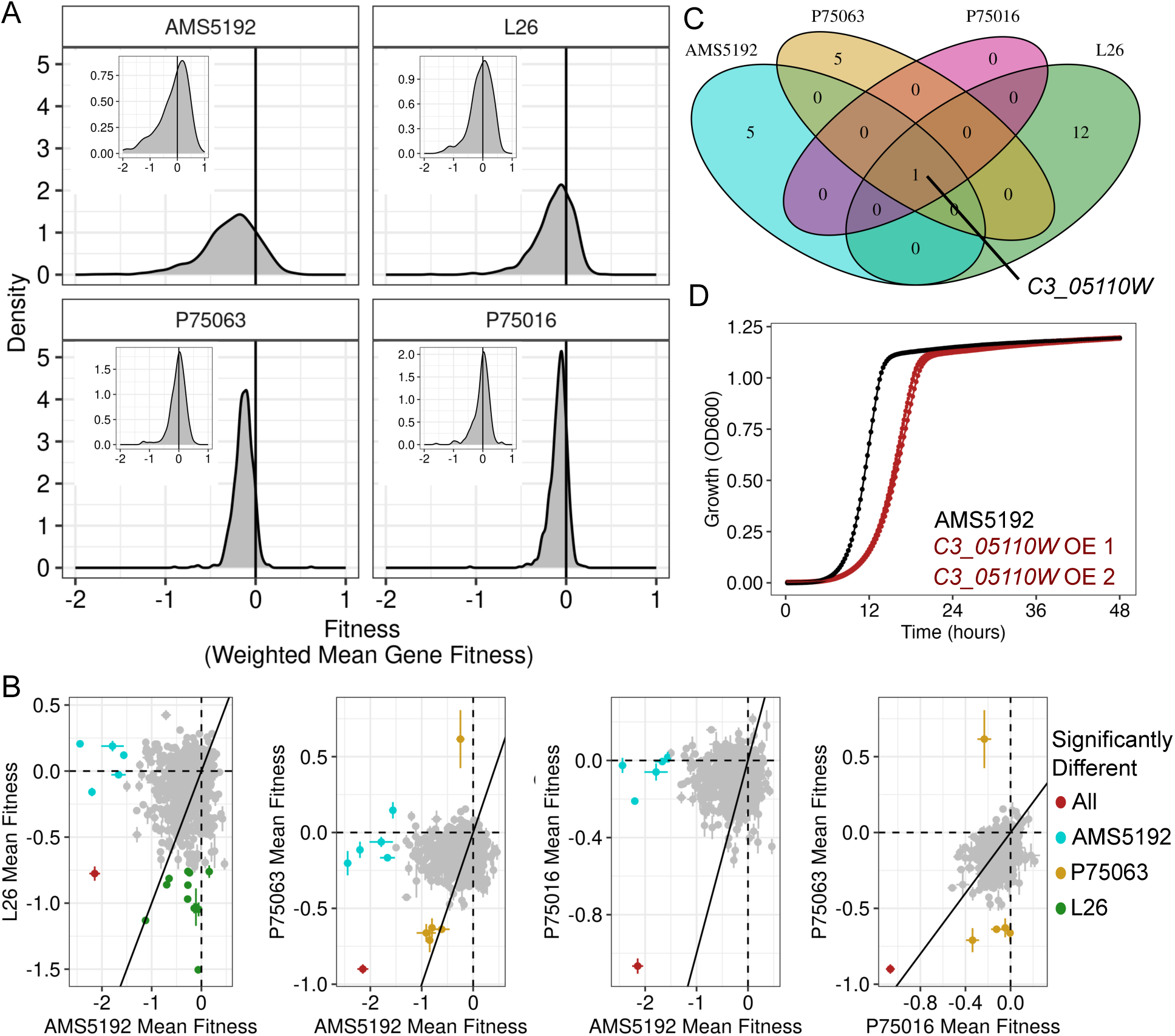
Fitness effects of gene overexpression in rich medium. (A) Density plots show the distribution of fitness effects of overexpression for all genes in rich medium for the four genetic backgrounds. Density plots for the same genetic backgrounds in rich medium for the non-targeting control strains are shown as insets in each panel. (B) Scatterplots show the fitness effects of overexpression for each gene (points) in one genetic background (x-axis) as compared to another genetic background (y-axis). Points are mean of three replicates and bars are standard errors of the mean. Genes with significant impacts on fitness are colored according to the strain in which they are significant (moderated t-test FDR < 0.05), or in red if they are significant in all backgrounds. (C) Venn Diagram showing intersection of genes with significant impacts on fitness across the four genetic backgrounds. (D) Growth measured as OD_600_ over time for the AMS5192 progenitor strain (black) and two independently generated transformants overexpressing *C3_05110W* (red). Points are mean of seven replicates and error bars are standard error of the mean.

### Fitness effects of gene overexpression are typically environment-specific

We next asked how the fitness effects of gene overexpression compare across different environments by identifying genes with significantly different fitness in each environment relative to the rich medium baseline (Fig. 3A, moderated t-test FDR <= 0.05, Supplementary Table S2). *CDR1* and *NCP1* in fluconazole had the largest effect sizes across all comparisons (Fig. 3A,D). These two genes have been previously associated with fluconazole resistance^36,39,45^ and our results indicate they are major drivers of CNV phenotypes in fluconazole.

**Fig. 3:**
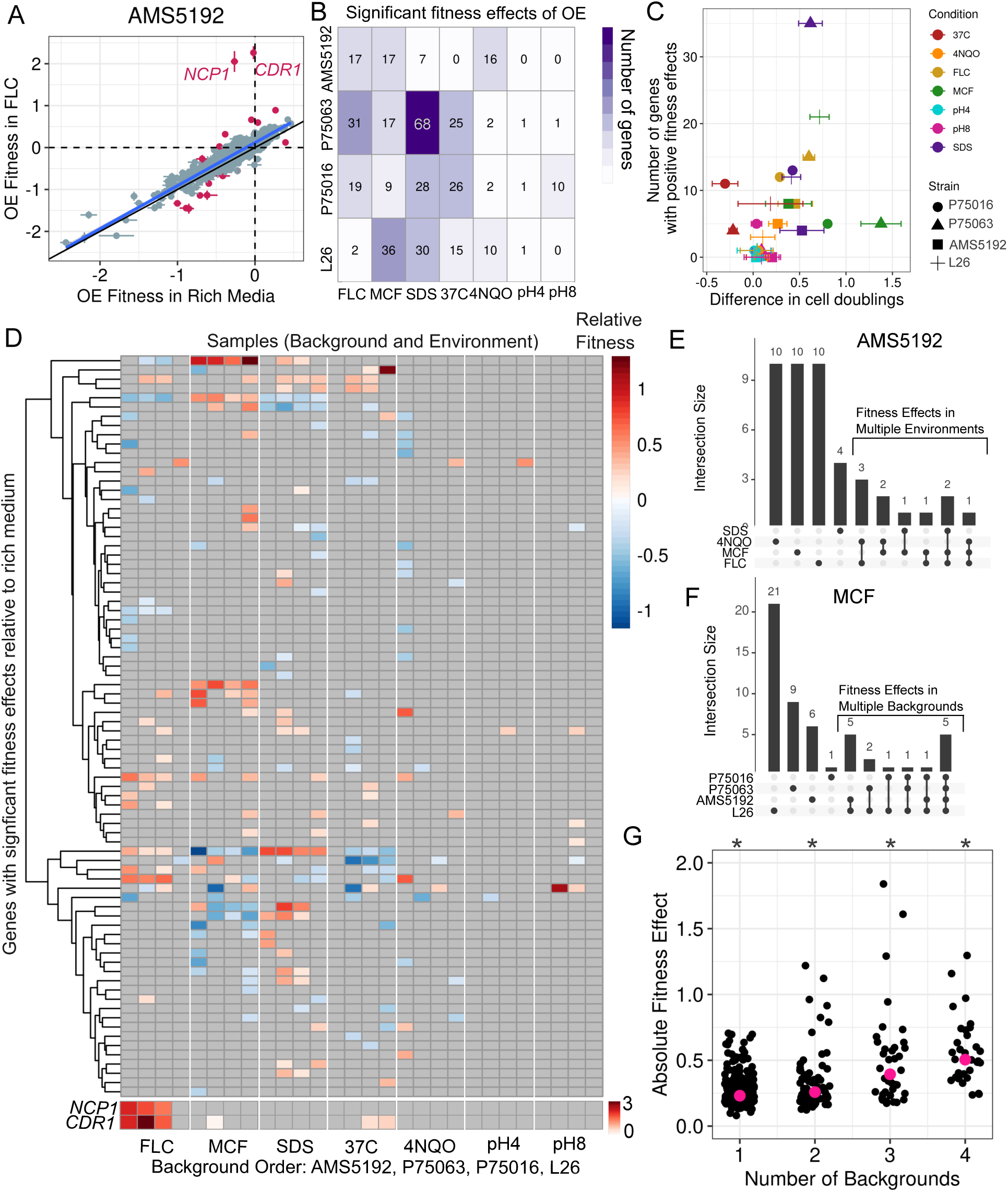
Fitness effects of individual gene overexpression across environments. (A) A scatterplot shows the fitness effects of overexpression for each gene (points) in rich media (x-axis) vs fluconazole (y-axis) for the genetic background AMS5192. Genes with significantly different fitness in fluconazole relative to rich media are colored in pink. Points are the mean of three replicates, error bars are standard errors of the mean, blue line is a best fit line, and black line is an x = y line. (B) A heatmap shows the number of genes with significant effects on fitness when overexpressed in each strain (rows) in each environment (columns). (C) The number of genes with significantly positive fitness effects in each environmental condition (colors) is plotted on the y-axis relative to the difference between the number of doublings in rich medium and that environment for each genetic background (symbols) on the x-axis (Linear model R^2^ = 0.148, p-value = 0.024). (D) A heat map showing the fitness effects of overexpression for genes (rows) with a significant effect on fitness (moderated t-test FDR <= 0.05 and log_2_ fold change >= 0.3) in each environmental condition and strain background (columns). Genes are clustered by hierarchical clustering as shown by the dendrogram on the left, and columns are not clustered but repeat the same background order across all 7 environments, shown at the bottom. *CDR1* and *NCP1* are shown separately for better visualization in the rest of the heatmap. (E) An upset plot shows all set intersections for genes with significant effects on fitness in SDS, 4NQO, MCF, and FLC in strain AMS5192. There were no genes with significant effects in the other environments in AMS5192. Intersection sizes are shown above for each environment with a darkened point, with bars connecting multiple environments for gene sets that affect fitness in both environments. (F) Upset plots show intersections for genes with significant effects in micafungin. As in E, intersection sizes are shown in the barchart above for each strain background with a darkened point below. Additional backgrounds and environments are shown in Extended Data Fig. 4. (G) The absolute fitness effect size of each gene overexpression is plotted on the y-axis binned according to the number of genetic backgrounds in which overexpression results in significant fitness effect in the same environment (x-axis). Large pink points are the median absolute fitness effect for all genes in the x-bin (ANOVA p-value = 8×10^−17^).

We identified a total of 198 genes with significant fitness effects in at least one environment in one or more genetic backgrounds. The genetic backgrounds differed in the number of genes with significant fitness effects in each environment (Fig. 3B). We asked whether there was a relationship between starting fitness of each strain in each environment and the number of genes that confer a positive fitness effect, consistent with diminishing-returns epistasis^19,44,46,47^ and found a weak correlation (Fig. 3C, R^2^ = 0.148, p-value = 0.024). This result indicates that while there may be some relationship between initial fitness of the progenitor and the fitness effects of gene overexpression, it does not completely explain the differences in number of significant fitness effects between genetic backgrounds.

In all four genetic backgrounds, most genes had significant fitness effects in only one environment (Fig. 3D,E, Extended Data Fig. 4). Similarly, across all environments most genes had significant effects on fitness when overexpressed in a single genetic background (Fig. 3D,F, Extended Data Fig. 4), indicating widespread genetic-by-environment interactions on the fitness effects of gene overexpression. Genes with significant fitness effects in multiple genetic backgrounds had on average larger effect sizes than those that were specific to one or two genetic backgrounds (Fig. 3G, ANOVA p-value = 8×10^−17^). This indicates that while there is an abundance of gene-by-environment interactions, gene overexpression with large fitness effects tends to be robust to genetic variation.

### Each CNV region contains a small number of genes with beneficial effects in fluconazole

We next looked for genes with positive fitness effects in fluconazole, the environment in which the CNVs evolved. We plotted fitness effects of overexpression in fluconazole relative to rich media for each gene according to its genomic position (Fig. 4A). The genetic background L26 has lower fluconazole sensitivity and a different set of genes with significant fitness effects compared to the other three backgrounds. Therefore, we focused on genes with significant effects in all three of the more sensitive strains. Remarkably, each chromosomal region included one gene with significant positive fitness effects in those three genetic backgrounds (Fig. 4A), including *TYE7*, *C3_03370C*, *CDR1*, and *NCP1*. The transcription factor *TYE7*, best known for regulation of the glycolytic pathway,^48^ has not been previously implicated in fluconazole sensitivity. We generated individual transformants overexpressing *TYE7* and performed a broth microdilution assay, which demonstrated that overexpression increases growth in fluconazole at the progenitor MIC_50_ across all backgrounds (Fig. 4B, two-sided t-test p-value < 0.05).

**Fig. 4:**
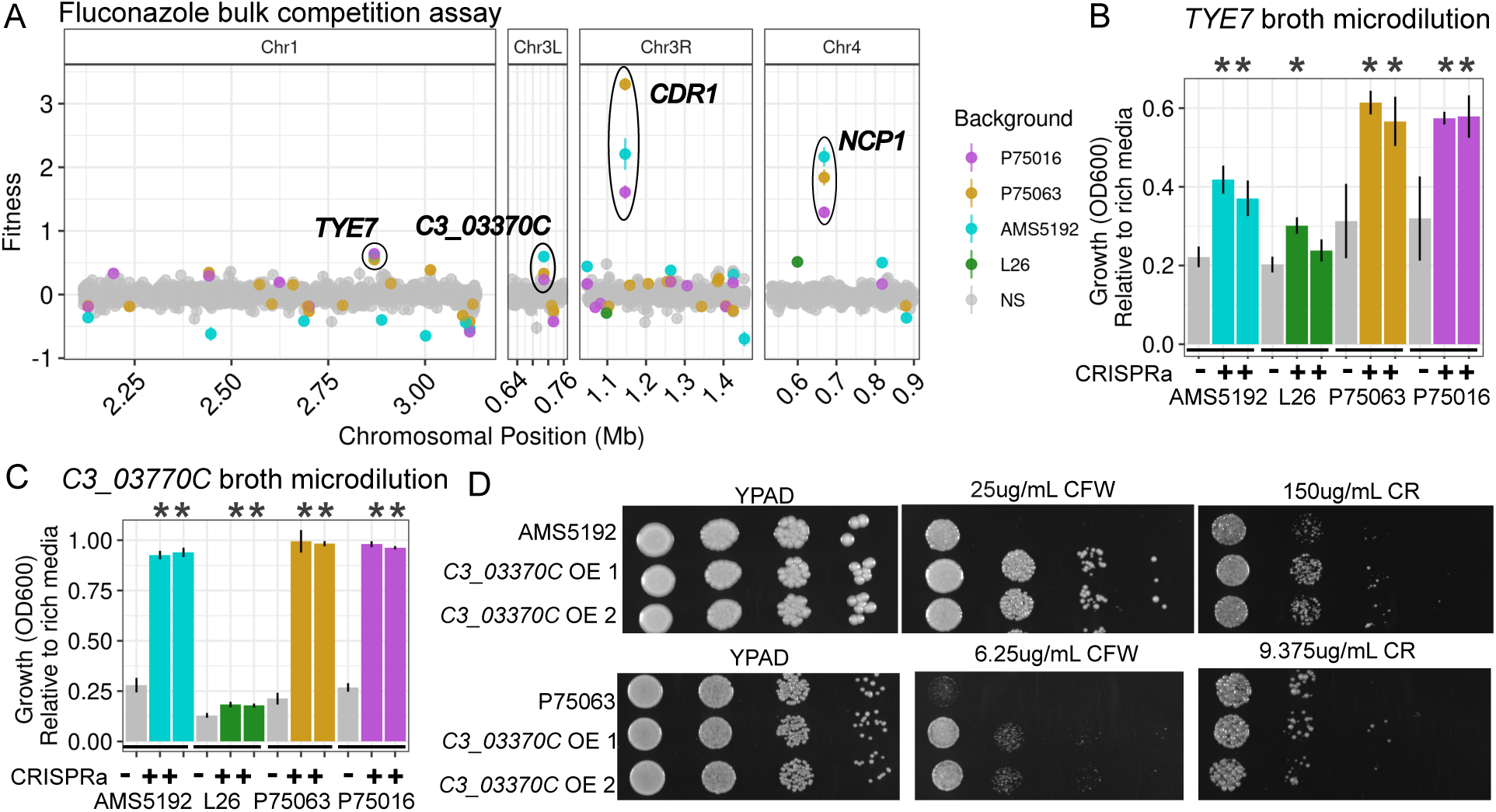
Overexpression of *TYE7* and *C3_03370C* increases fitness in fluconazole. (A) Fitness values in fluconazole relative to fitness in rich media are plotted on the y-axis, relative to the genomic position of each gene for regions of chromosomes 1, 3L, 3R, and 4 targeted by CRISPRa. Each point is mean fitness across replicates for one gene and bars are standard error of the mean. Genes with significantly different fitness in fluconazole are colored by the genetic background in which they are significant. Ellipses circle genes with significantly increased fitness in fluconazole when overexpressed in 3 genetic backgrounds and labeled. (B) Growth at the progenitor’s fluconazole MIC_50_ (1µg/mL, 4µg/mL, 1µg/mL and 1µg/mL FLC, respectively) relative to growth in rich media for each progenitor (grey bars) and two independently engineered transformants overexpressing *TYE7* (colored according to genetic background as in panel A). Bars are the mean relative growth and error bars are the standard deviation of the mean. Asterisks denote CRISPRa transformants with significantly different growth than their matched progenitor strain (two-sided t-test p-value < 0.05) (C) As in (B), relative growth in fluconazole of independently generated mutants overexpressing *C3_03370C* are plotted for each genetic background. (D) Cells spotted onto agar containing either YPAD media with no additives, calcofluor white (CFW), or Congo red (CR) from left to right are shown after 24 (CFW) or 48 (CR) hours of growth. Spots are at cell densities of 10^5^, 10^4^, 10^3^, and 10^2^ cells/mL from left to right. Progenitor strains (top rows) are compared to two independently generated mutants overexpressing *C3_03370C* (middle and bottom rows). See Extended Data Fig. 7 for additional genetic backgrounds.

The uncharacterized gene *C3_03370C* has also not been previously associated with fluconazole sensitivity. Independent transformants overexpressing *C3_03370C* in each background exhibited a 2-fold increase in fluconazole MIC_50_ in the three sensitive backgrounds and slightly increased growth in L26 at the MIC_50_ (Fig. 4C, two-sided t-test p-value < 0.05, Extended Data Fig. 5). To probe possible roles of this protein and the mechanism underlying fluconazole sensitivity, we tested additional antifungals and cell wall stress with our individually generated mutants. We did not observe a change in micafungin (Extended Data Fig. 6) but did see decreased sensitivity to Calcofluor White and Congo Red (Fig. 4D, Extended Data Fig. 7). Together these results suggest that overexpression of *C3_03370C* influences cell wall integrity, which might be related to the mechanism of fluconazole susceptibility.

### Overexpression of several genes exhibit fitness trade-offs across environments

CNV regions also include genes with significant fitness effects in other environments. In micafungin, a second class of antifungal drug with a distinct mechanism of action, we found that overexpression of *C1_11720W, SET3*, *C3_03460C*, *YCK2*, and *SKN1* significantly affected fitness in all four genetic backgrounds (Fig. 5A, moderated t-test FDR <= 0.05). None of these five genes have been previously associated with sensitivity to micafungin. Three of these genes, *C3_03460C, SET3,* and *YCK2*, had also had significant effects in fluconazole (Fig. 5B, moderated t-test FDR <= 0.05), and in SDS in multiple backgrounds (Fig. 5C, moderated t-test FDR <= 0.05). Broth microdilution assays for independent transformants recapitulated all phenotypes observed in the bulk competition assay (Fig. 5D, Supplementary Table S3). We asked whether there were general trade-offs between fitness effects in fluconazole, micafungin, and SDS. There was no relationship between micafungin and fluconazole (Fig. 5E, linear model AdjR^2^ = −0.009, p-value = 0.368) but there was a significant negative relationship between micafungin and SDS for genes with significant effects in both environments (Fig. 5F, linear model AdjR^2^ = 0.31, p-value = 0.0015). *C3_03460C* showed a unique pattern of trade-offs across the three environments as compared to *YCK2* and *SET3* (Fig. 5G), indicating that while all three genes affect fitness in all three environments, the nature of these trade-offs are gene specific and suggest different mechanisms of action.

**Fig. 5:**
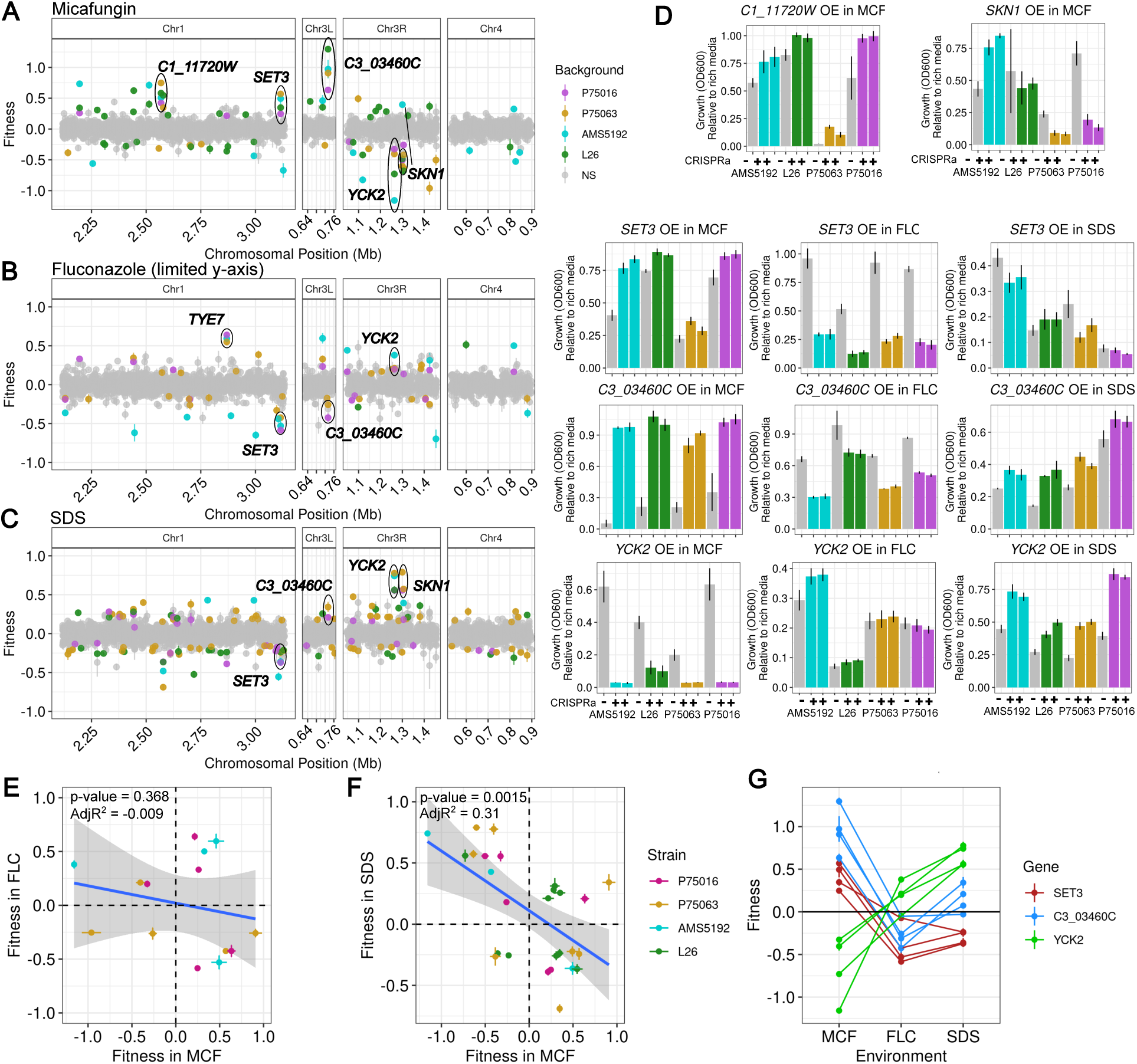
Fluconazole-evolved CNVs contain genes whose overexpression affects fitness in micafungin and SDS. (A-C) The fitness effects of overexpression of individual genes in micafungin (A), fluconazole (B), and SDS (C) relative to rich media are shown according to their genomic position for ranges of chromosomes 1, 3L, 3R, and 4 targeted in the CRISPRa pooled assay. The fluconazole data is the same as Fig. 4A, but with a limit on the y-axis removing *CDR1* and *NCP1* to better visualize the rest of the data. Points are the mean of three replicates and bars are standard error of the mean. Each point is colored grey if not significant in any genetic background or colored according to the genetic background where it is significant. (D) Growth as measured by OD_600_ in micafungin, fluconazole, or SDS relative to growth in rich media for each genetic background (grey bars) and two independently generated overexpression mutants in each of the four genetic backgrounds (colored according to background as in panel A) are shown. Bars are the mean relative growth and error bars are the standard deviation. Micafungin concentration is 0.016 ug/mL, fluconazole concentrations are at the progenitor MIC_50_ or 2-fold below the progenitor MIC_50_ for decreases in fitness, and SDS concentrations are at the progenitor MIC_50_ (see Methods, Supplementary Table S3). (E) For genes with significant fitness effects in both micafungin and fluconazole, the fitness effect observed in micafungin is plotted on the x-axis, and the fitness effect in fluconazole is on the y-axis. Points are colored by the genetic background in which they are significant in both environments. Points are means and error bars are standard error of the mean across three replicates. The blue line is a best fit line and grey shaded areas are 95% confidence intervals. (F) As in (E), but for genes with significant fitness effects in both micafungin (x-axis) and SDS (y-axis). (G) Fitness values for overexpression of *SET3* (red), *C3_03460C* (blue), and *YCK2* (green) are connected via line segments across environments (x-axis) for all four genetic backgrounds. *SET3* and *YCK2* show trade-offs between micafungin and both fluconazole and SDS, while *C3_03460C* shows a trade-off between micafungin and fluconazole but not SDS.

### Single gene overexpression can explain fitness trade-o>s between environments for CNVs

By amplifying many genes, CNVs might result in fitness trade-offs between environments that could be exploited to combat adaptation to drug.^49^ However, how the fitness effects of individual gene amplifications relate to the fitness effects of an entire CNV across environments is unknown. Therefore, we examined how well a simple additive model for individual gene overexpression could predict the fitness of a strain bearing a complex CNV. Strain P75016-Chr3R-CNV contains a single complex CNV on chromosome 3R (Fig. 6A) that includes *CDR1, YCK2*, and *SKN1* amongst 139 other genes, several of which have significant fitness effects in different environments (Fig. 6B). We performed growth curve assays for P75016-Chr3R-CNV in each environmental condition and calculated its growth relative to its euploid progenitor P75016. Strikingly, the additive fitness effects from individual gene overexpression could significantly predict the CNV effect on growth (Fig. 6C, linear model AdjR^2^ = 0.36, p-value = 0.002), primarily driven by increased fitness in fluconazole and decreased fitness in micafungin. The same trend was observed for two different genetic backgrounds with different CNVs on chromosome 3R (P75063-Chr3R-CNV, Extended Data Fig. 8) and chromosome 4 (SC5314-Chr4-CNV, Fig. 6D-F). In contrast, one isolate with the largest CNV on chromosome 1, P75063-Chr1-CNV, had increased growth in micafungin as expected from the single gene overexpression effects, but increased growth in fluconazole despite additive single gene fitness data being negative (Fig. 6G-I). The incongruence of the fluconazole phenotype for P75063-Chr1-CNV suggests that other factors such as epistatic effects between amplified genes or other structural variant effects might be at play in this isolate, in contrast to the previous isolates that are described well by individual gene fitness effects.

**Fig. 6:**
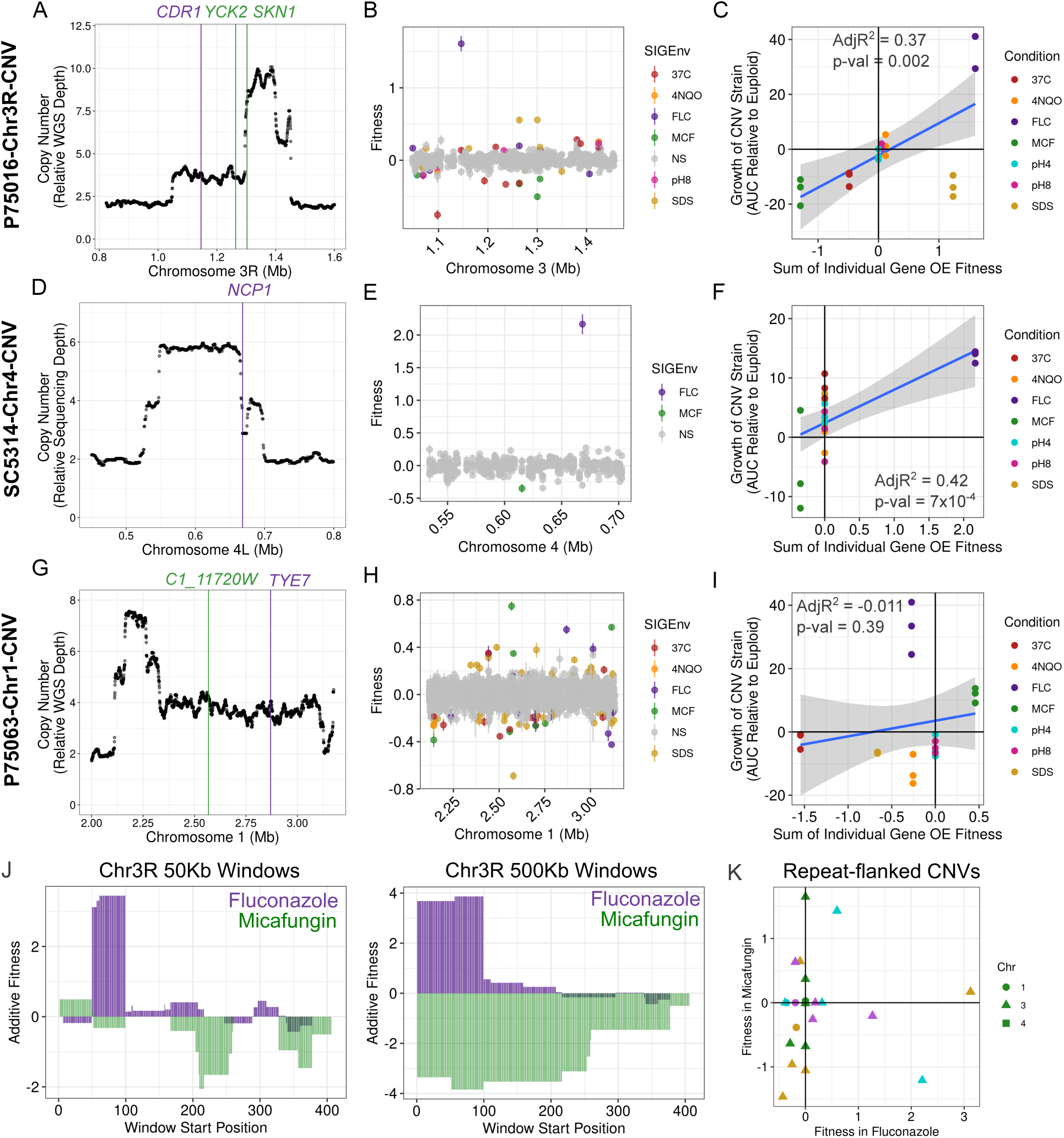
Individual gene overexpression fitness predicts CNV growth and fitness trade-offs. (A) A rolling median of mean whole genome sequencing read depth normalized to median read depth is plotted on the y-axis across the genomic positions shown on the x-axis for a region of chromosome 3R in strain P75016-Chr3R-CNV. Vertical lines show the positions of the genes encoding *CDR1*, *YCK2*, and *SKN1*. (B) Fitness of single gene overexpression from the CRISPRa bulk competition assay is shown for the region of chromosome 3R amplified in strain P75016-Chr3R-CNV. Each gene is shown multiple times for each environmental condition, but only background P75016 is shown. Genes without significant fitness effects are shown in grey, while genes with significant fitness effects are colored according to the environment in which they have the effect. Points are means of three replicates and bars are standard error of the mean. (C) Relative growth, as measured by the area under the curve (AUC) of strain P75016-Chr3R-CNV relative to its matched progenitor strain, P75016, is plotted on the y-axis. Growth in each condition relative to rich media is shown in different colors as indicated in the legend.

Triplicate measures are shown as individual points. Predicted fitness effect by summing the significant fitness effects of all genes in the amplified region in P75016-Chr3R-CNV are plotted on the x-axis. Because there is only one predicted fitness by sum, but replicate measures for the CNV strain, there is one x-axis value for each of three y-axis measures. Replicate measures of CNV growth are shown separately due to potential instability of the CNV during growth. (D-I) As in A, B, and C, but for strain SC5314-Chr4-CNV, in which a portion of chromosome 4 is amplified and is compared with SC5314-derived background AMS5192, and P75063-Chr1-CNV, in which a portion of chromosome 1 is amplified and is paired with background P75063. (J) Predicted fitness effect of 50Kb windows (left) or 500Kb windows (right) paneling across the region of chromosome 3R by adding individual gene fitness are shown for fluconazole (purple) and micafungin (green). (K) Fitness in fluconazole (x-axis) and micafungin (y-axis) for all potential CNV regions flanked by inverted repeats within the CNV regions assayed by CRISPRa. Points are shaped according to which chromosome the CNV would be located on and colored by genetic background.

We hypothesized that the precise boundaries of a CNV would have a significant impact on fitness trade-offs because of the different genes contributing to fitness in each environment. We calculated the additive fitness effects of gene overexpression for 50Kb and 500Kb windows scanning across chromosome 3R at 1Kb intervals in fluconazole and in micafungin for background P75063 (Fig. 6J, Extended Data Fig. 9). Trade-offs differed depending on boundaries and larger CNVs showed larger and more consistent fitness trade-offs (Fig. 6J, Extended Data Fig. 9). These analyses support the hypothesis that larger CNVs include individual genes with positive and negative impacts on fitness in different environments, resulting in fitness trade-offs.

CNVs in *C. albicans* are flanked by long inverted repeats that play an important role in their formation.^20,37,38^ We asked whether CNVs could be refined using existing inverted repeat sequences within regions included in our bulk competition assay. We calculated additive fitness effects for all possible regions flanked by inverted repeats in fluconazole and micafungin and found that only two regions resulted in high fitness in fluconazole without a fitness cost in micafungin (Fig. 6K). These data suggest that smaller CNVs reduce fitness trade-offs between classes of antifungal drug, however the position of long inverted repeats limit further re-sizing.

## DISCUSSION

Our systematic CRISPR-activation screening data provide the first comprehensive view of the contributions of individual genes to the fitness effects of complex CNVs in *C. albicans,* including how they vary across diverse genetic backgrounds and environments, with important implications for how CNVs might be expected to contribute to adaptation in complex environments such as the human host. We identified several novel genes involved in antifungal drug susceptibility. Below we highlight how this unique and comprehensive dataset reveals several key findings that help explain the fitness effects of CNVs.

First, many fitness effects of gene overexpression show gene-by-environment interactions. These findings lend additional support to the move toward testing gene function in multiple genetic backgrounds rather than relying on a single reference strain.^19,23,50^ For example, overexpression of *SKN1* results in an opposite direction of effect in micafungin in AMS5192, a lab reference strain, as compared to the other three backgrounds (Fig. 5D), despite being similarly overexpressed by CRISPR-activation (Extended Data Fig. 5). This difference in the reference strain may help explain why *SKN1* has not been previously associated with drug sensitivity phenotypes. More work is needed to better understand the basis of genetic background effects on a genome-wide scale. However, we also find that the fitness effects of overexpression that are consistent across genetic backgrounds are also those with the largest effect sizes. This is encouraging, because both large effect sizes and consistency across backgrounds are desirable for the selection of potential drug targets.

Second, we systematically identified individual genes that contribute to the fitness gains in each complex CNV strain in antifungal drug when overexpressed, most of which have not previously been tied to drug sensitivity phenotypes. Most functional gene annotation is based on loss-of-function phenotypes, and while these are productive approaches, the effects of gene overexpression are not well predicted by loss-of-function mutants. *C3_03370C* is a short peptide that has not been previously characterized in *C. albicans* and does not have an ortholog outside of *Candida*. Overexpression of this gene also influences sensitivity to Calcofluor White and Congo Red, suggesting it plays a role in cell wall integrity. Further work is needed to elucidate the mechanism by which it influences azole sensitivity. The specificity of gene overexpression fitness effects might also impact their involvement in adaptation to antifungal drug. For example, *CDR1* overexpression has a strong positive fitness benefit in fluconazole without significant negative fitness effects in any other environment tested. The lack of negative consequences for *CDR1* overexpression might help explain its prominence as a mechanism of drug resistance across diverse clinical isolates.^45,51–53^

Third, CNVs amplify a small number of genes with positive fitness effects in the selection environment along with additional genes that can have negative fitness consequences elsewhere. Relatively few genes appear to be the drivers of decreased fluconazole susceptibility within each CNV. The amplification of what appears to be unnecessarily large regions is likely due to mutational mechanisms that allow these regions to be amplified at high frequencies.^5,21,36,38,54^ The amplification of many additional genes does not negatively affect fitness in fluconazole, the selection environment, but does include genes with negative fitness effects in micafungin. Our simulated CNVs predict that CNVs affecting fewer genes would be less likely to have negative fitness trade-offs in micafungin. However, there are very few inverted repeat pairs within these regions that would still result in amplification of the driver genes. This suggests that it may be difficult to refine a CNV in *C. albicans*, leaving strains bearing these CNVs vulnerable to micafungin therapy.

The development of drug resistance via CNVs is a major concern for fungal pathogens where treatment options are limited. The trade-offs we observe here are encouraging for the potential of combination or step-down therapies that employ multiple drug classes to treat isolates adapting to drug via CNVs. Future studies examining the fitness effects of gene overexpression genome-wide will show how well these recurrently amplified regions represent the genome as a whole.

## METHODS

### Generation of fungal CRISPR-activation pools

#### Development of guide pools

Guides were designed as described previously^40^ and a detailed protocol for guide design and library build can be found on Protocols.io^55^ and https://github.com/TheShapiroLab/Pooled-Library-Build/tree/main. A list of all features encoded on Chr1: 2117999-3133121, Chr3: 621867-897198 and 1047995-1457807, and Chr4: 531821-898541, corresponding to regions amplified in CNVs,^20,37,38^ was extracted from the feature file C_albicans_SC5314_version_A21-s02-m08-r15_chromosomal_feature.tab downloaded from the Candida Genome Database (candidagenome.org^56^) and names converted to *Candida albicans* assembly 22 systematic names (Supplementary Table S4). All sequence information for each of these 843 features was downloaded from the Candida Genome Database using SC5314 assembly 22. Sequence ranges, as determined below, were isolated and inserted into the guide design tool located at http://grna.ctegd.uga.edu. Ranges targeted were −90bp to −370bp upstream of only the start codon of ORF features that had a predicted dominant TSS of less than or equal to 140bp, or of upstream of both the TSS and the start codon for ORF features with a predicted dominant TSS of greater than 140bp. If features did not have a predicted dominant TSS, ranges −90bp to -670bp were targeted upstream of the start codon. For non-ORFs (tRNAs, long terminal repeats, snoRNAs, retrotransposons, snRNAs, blocked reading frames, pseudogenes, and direct repeats) ranges +100bp to −400bp from the feature start site were targeted.

From the list of sgRNAs generated by the online tool, all sgRNAs with any predicted perfect or imperfect off-target effects were removed, as well as any sgRNAs with “N”s in their sequence. In addition, all guides that contained a Pac1 or Sap1 restriction site were removed to avoid issues when cloning. From this list, the following numbers of highest-scoring guides for each feature type were kept: 8 sgRNA for ORF features that had a predicted dominant transcription start site of less than or equal to 140bp; 4 sgRNAs targeting upstream the start codon and 5 sgRNAs targeting the TSS for ORF features that had predicted dominant TSS of greater than 140bp; 5 sgRNAs within 370bp of the start codon and 4 sgRNAs greater than 370bp from the start codon for ORF features with no predicted dominant TSS; and up to 10 sgRNAs for non-ORF features. In addition, 60 sgRNAs that have no predicted homology of greater than 15bp in the *C. albicans* assembly 22 genome as determined by the Candida Genome Database BLASTn tool were added to constitute ∼1% of the pool. Flanking sequences for golden gate assembly were added to each guide sequence, and this final list of oligos was synthesized in pooled oligo format by Twist Biosciences (Supplementary Table S4).

#### Generation of bacterial pool

For golden gate cloning of the oligo pool into the CRISPR-activation plasmid, plasmid pRS156^39^ (AddGene #182707) was mini-prepped using a Qiagen Mini-Prep Kit. The Twist Oligo Pool was centrifuged at full speed for 10 seconds and resuspended in TE (pH 8.0) to 20 ng/µL at 55°C by pipetting. PCR using KAPA HiFi polymerase was then performed for 10 cycles to amplify the pool using primers 2302 and 2303 (Supplementary Table S5). After amplification, the pool was cleaned up using a commercial PCR clean up kit (Zymo Research Clean and Concentrator Kit). As previously described for pooled CRISPRi assays^50^, a golden gate reaction was then performed using a 1:3 vector:insert ratio, with a total of ∼5000 ng of vector. The golden gate reaction product was then transformed into NEB 10-beta *E. coli* electrocompetent cells by electroporation, and cells were incubated in semi-solid agar (SeaPrep) LB media containing 100 µg/mL ampicillin and 250 µg/mL nourseothricin. Colonies were allowed to grow at 30°C for two days before being centrifuged, washed, and glycerol stocked. A maxi-prep (NucleoBond Xtra Maxi EF) was then performed with the bacterial pool according to the manufacturer’s instructions.

#### Generation of the fungal pools

Fungal pools were generated by first maxi-prepping the bacterial plasmid pool (NucleoBond Xtra Maxi EF) and digesting with Pac1 at 37°C for 12 hours and then heat-inactivating at 65°C for 20 minutes. One 50 µL digestion of ∼5000 ng of plasmid pool was prepared for each yeast transformation and 25 yeast transformations plus one negative control transformation was performed for each yeast genetic background. For each yeast genetic background, a small amount of the background strain was patched onto YPAD agar (20 g/L Bactopeptone, 10 g/L yeast extract, 0.04 g/L adenine, 0.08 g/L uridine, 2% dextrose) and grown for ∼2 days at 30°C. A small dot was used to inoculate 50 mL of YPAD liquid medium and grown overnight at 25°C. Provided the OD_600_ was close to 5, and not below 2 or above 7, the following equation was used to calculate the volume of the overnight culture to use for each transformation: volume to use = 10,000/(OD_600_x1.5). This volume was then spun down in a microcentrifuge tube and the supernatant decanted for each of the 26 transformations. Each transformation was then resuspended in 1 mL of a transformation mix (800 µL 50% PEG, 100 µL TE, 100 µL 1M Lithium acetate, 20 µL boiled sheared salmon sperm DNA, 20 µL DTT per transformation). 50 µL of the overnight Pac1 digested plasmid pool was then added to each transformation and inverted several times to mix. Cells were then incubated at 30°C for 1 hour and then subjected to heat shock at 42°C for 45 minutes. Each transformation was then spun down at 400xg for 5 minutes and the transformation mix pipetted off. Cells were washed in 1 mL YPAD 3 times. After the final resuspension, all 25 positive transformations were added to a 500 mL Erlenmeyer flask containing 50 mL of YPAD and shaken gently at 93 rpm at 30°C for 3 hours. All 75 mL were then gently spun down and resuspended in 2.5 mL of YPAD and carefully mixed. 150 µL aliquots were then plated on YPAD agar containing 100 µg/mL nourseothricin and 1 mg/mL quinine and spread using glass beads. In addition, two 10x dilutions of the resuspended cells were plated for counting. The negative control was plated one on plate at the full concentration. Plated cells were allowed to grow at 30°C for 2 days.

After 2 days of growth, individual colonies were visible on both full concentration and 10x dilution plates, while the negative control plates remained clear. Cells were counted on 10x dilution plates and individual colonies patched for gDNA extraction and PCR. PCR was used to check for correct insertion of the CRISPRa construct (primers 1744 and 1745, Supplementary Table S5) and the sgRNA sequence amplified (primers 1743 and 1742, Supplementary Table S5) and sent for Sanger sequencing to ensure unique guides present in each colony tested. Full concentration transformation plates were scraped and cells were resuspended in YPAD. After thorough mixing and resuspension in 25 mL YPAD, pools were glycerol stocked in 1 mL aliquots in 20% glycerol.

#### Fungal pool bulk competition assays

For each of three replicate assays, one aliquoted glycerol stock of each fungal pool in the four genetic backgrounds was defrosted on ice and a small amount used to make a 10x dilution and measure the OD. Defrosted glycerol stocks were back diluted to an OD_600_ of 0.5 in 10 mL of YPAD. Cultures were incubated at 30°C with shaking for 5 hours. After this period the OD_600_ of a 10x dilution was again measured to ensure that it had reached on OD_600_ of ∼0.5. 1.5 mL of each culture was then pelleted and frozen in liquid nitrogen. Each culture was then back diluted to reach a final OD of 0.05 in 10 mL of each environmental condition media. Cultures were placed on a rotating wheel at 30°C (or in a shaker at 37°C for the 37°C condition) and grown for 24 hours. After 24 hours, OD_600_ was measured and 1.5 mL was removed from the culture, pelleted, supernatant removed, and frozen in liquid nitrogen. Changes in OD_600_ from initial timepoint to 24 hour timepoint was used to estimate number of doublings for each sample. At 36 hours this procedure was repeated for a second timepoint. Upon analysis, the first and second timepoints were very similar in their fitness values and therefore only the first timepoint was used for simplicity.

Genomic DNA was extracted from all frozen cell pellets as described in methods section ‘gDNA extraction’. DNA concentrations were normalized to ∼5 ng/µL and used as input to a low-cycle PCR using primers 2010 and 2011 to amplify the region of the genome in which the sgRNA is integrated and the PCR product cleaned using Ampure XP (Fisher Scientific A63881) magnetic beads at 1.8x the reaction volume. A second low-cycle PCR was then performed on the cleaned product to add Illumina indexed adapters with sample-specific dual indexes (Illumina DNA/RNA Unique Dual Indexes Set A, Set B, and Set C), and the PCR product once again cleaned using Ampure XP magnetic beads. The concentration of each sample was then quantified using Qubit 4 Fluorometer (Invitrogen) dsDNA high sensitivity kit. Concentrations were used to calculate the volumes of each sample needed to normalize all concentrations to 10 ng/µL when pooled. All samples were then pooled, the final concentration verified using Qubit at 9.53 ng/µL and submitted to University of Minnesota Genomics Center for sequencing. Pooled samples were sequenced on the Element AVITI Freestyle 2×150 High run (1000 M read output).

### Statistical Analysis of pooled competition assays

#### Quantification of barcodes at each timepoint

Reads for each sample were de-multiplexed by the University of Minnesota Genomics Core and fastq files for each sample delivered to the Selmecki Lab. As described previously,^40,50,55^ read pairs were first assembled using Pear^57^ (v0.9.11) with maximum overlap of 200 and minimum overlap of 100. 50 bp were trimmed from the ends of merged reads and remaining read sequence reverse complemented using vsearch (v2.3.4). Reads with perfect matches to each sgRNA sequence in the library plus an 8 nucleotide anchor on each side were then counted using a parallelized^58^ “zgrep -c” command on the University of Minnesota Supercomputing Institute computing cluster. These final barcode counts for each sample appear in the data matrix “Allrawcounts.txt” (Supplementary Table S6) and all code used for processing is available at https://github.com/selmeckilab/CRISPRactivation_for_CNV_genes.

#### Calculation of gene-level fitness values

Statistical analysis of barcode counts to generate fitness estimates for each gene in each sample were conducted in R (v4.4.0) using an edited script for estimating fitness of Tn-seq bacterial pools from Martinson et al, 2023.^59^ The edited script used for analysis is available at https://github.com/selmeckilab/CRISPRactivation_for_CNV_genes. Barcodes with fewer than 50 reads present in initial timepoints and samples with less than a median of 50 reads per barcode were removed from analysis. Genes that were then targeted by only one barcode were removed from analysis. Each sample was normalized using the depth of the reference (non-targeting) barcodes. Fitness of each barcoded strain was calculated as the log_2_ fold change in barcode frequency from the sample-specific initial timepoint to the 24 hour timepoint for each pool and environmental condition. We found that barcoded strains with high variance related to low initial frequencies tended to have individual strain fitness values that deviated strongly from other sgRNAs targeting the same gene (Extended Data Fig. 3). For this reason, individual barcode fitness values were weighted by multiplying by the inverse of their variance to penalize barcoded strains with high variances, with a weight cap set at the variance value of a barcode with an initial frequency of 20. These weighted individual barcode fitness values still do not show uniform fitness effects across all guides targeting the same gene, and are not expected to, as individual guides may induce overexpression to different degrees and in some cases even interfere with gene transcription and result in repression rather than overexpression (Extended Data Fig. 3). We reasoned that these unintended effects would be less frequent and less consistent across sgRNAs than the effects of inducing overexpression as intended and therefore calculated an average fitness effect of all sgRNA barcoded strains targeting the same gene to produce a gene-level fitness value for each sample. These composite fitness values are a conservative estimate of gene overexpression fitness and also reduce the multiple testing burden. In addition, individually regenerated mutants validated that the weighted average gene-level fitness values are representative of gene overexpression (Figures 1,4,5, Extended Data Fig. 5). Finally, fitness values were adjusted so that the median fitness of the non-targeting control barcodes was zero (these adjustments were uniformly small, as median fitness for non-targeting controls were already close to zero). All statistical analyses described in the main text use these gene-level weighted average fitness values across three replicate competitions for each strain background and environmental condition.

#### Additional QC for between-pool comparisons

When making comparisons between genetic backgrounds the exact composition of the pool is confounded with the genetic background because each background pool was generated independently. To test how much the exact composition of the pool might influence fitness values, we generated two independent pools in the same background, P75063, and conducted a competition experiment in rich media for both independently generated pools side-by-side (P75063A and P75063B). The distribution of fitness effects in rich media were very similar between P75063A and P75063B (Extended Data Fig. 10A,B), and genes with large fitness effects were generally identical between pools (Extended Data Fig. 10C). However, there were also genes whose fitness values in rich media differed between P75063A and P75063B (Extended Data Fig. 10C). Therefore, we were careful to perform all between-pool comparisons after first normalizing to a baseline condition and therefore accounting for pool-specific differences in barcode frequency changes.

### gDNA extraction

gDNA extraction for barcode sequencing of bulk competitions and for whole genome sequencing of CNV strains and independently generated CRISPRa transformants was performed using the same protocol. Cells were pelleted and resuspended in TENTS buffer (1% SDS, 100 mM NaCl, 100 mM Tris pH 8, 1 mM EDTA, 2% Triton). ∼250 µL of .5 mm glass beads was added and cells were lysed in a BeadRuptor Elite (Omni International, 1 cycle, 4 m/s, 15 s). Genomic DNA was then isolated using a phenol:chloroform extraction and resuspended in DEPC-treated water.

### Whole genome sequencing of CNV single colonies

As described previously^36^ sequencing libraries for whole genome sequencing were prepared by SeqCenter, LLC, using the Illumina DNA Prep and sequenced on an Illumina NovaSeq 6000. Adaptor sequences and low-quality reads were removed using Trimmomatic (v0.39, parameters LEADING:3 TRAILING:3 SLIDINGWINDOW:4:15 MINLEN:36 TOPHRED33).^60^ Trimmed reads were mapped to the *C. albicans* A21 reference genome (A21-s02-m09-r08) from the Candida Genome Database.^56^ Reads were mapped using BWA-MEM (v0.7.17) with default parameters.^61^ Mapped reads were then sorted, indexed, and PCR duplicates removed using Samtools (v1.10)^62^ and read depth at each genomic nucleotide position calculated using the ‘samtools depth’ function.

For plotting of CNVs in single colonies isolated from evolved CNV populations, depth files were further processed in R (v4.4.0) to identify specific CNVs. Depth files were read into R, and rolling means were calculated for 500 bp windows across the entire genome using the RcppRoll R package (v0.3.0).^63^ Means were then divided by the median read depth of the entire genome (excluding mitochondrial DNA) to calculate relative abundance and then multiplied by a ploidy of 2 to generate an estimated copy number. For final smoothing of these copy number estimates, a rolling median of 25 means was calculated and plotted according to genome position to visualize CNVs. Long inverted repeat positions were downloaded from Supplementary File 2 from Todd et al, (2019),^38^ and major repeat sequences added from the Candida Genome Database.

### Generation of individual overexpression mutants

For independent generation of transformants overexpressing genes of interest from the pooled assays, one sgRNA sequence was selected from all guides targeting a gene of interest that most closely matched the weighted average fitness effects of all guides targeting that gene. This guide sequence with flanking regions for golden gate cloning was then ordered in forward and reverse orientation (primers for each strain are listed in Supplementary Table S5) from IDT. Oligos were resuspended in duplex buffer (30 mM HEPES, pH 7.5; 100 mM Potassium acetate, filter sterilized) and duplexed by first heating individual tubes to 94°C for 1 minute, adding equal volumes of each primer to a new microcentrifuge tube, letting the mixture sit for 1 min, heating to 94°C for 2 minutes, and finally returning to room temperature. Guides were inserted into the digested CRISPRa vector (pRS156, Supplementary Table S5)^39^ using the same golden gate method as for the pooled assays, transformed into NEB 5-alpha *E. coli* cells using heat shock, and positive colonies selected for on LB agar containing 100 µg/mL ampicillin and 100 µg/mL nourseothricin after growth at 30°C for 2 days. Single colonies were checked for the correct plasmid by PCR using the guide primer and primer 1742 (Supplementary Table S5). Correct colonies were then grown in liquid LB containing 100 µg/mL ampicillin, glycerol stocked, and the plasmid mini-prepped (Qiagen QIAprep Spin Mini Prep Kit).

Plasmid was digested overnight with Pac1 restriction enzyme and transformed into each yeast genetic background using the same protocol as used for the bulk transformations described above, but with only one transformation for each genetic background. Single colonies growing on selective plates were tested by PCR using the same primers as described above and Sanger sequencing to confirm the present of the correct guide. A minimum of 3 colonies that were confirmed by PCR were glycerol stocked for each individual mutant and 2 were subsequently phenotyped. If the two mutants showed different phenotypes, the third mutant was tested and the two mutants in agreement were presented in the results.

### Broth microdilution and growth curve assays

For broth microdilution assays and growth curve assays, strains were grown from frozen glycerol stocks overnight in 3 mL of YPAD. Strains were diluted to an OD of 0.01 in YPAD containing 1% glucose and used to inoculate the assay plate at 1:10, resulting in a final starting OD of 0.001 in the assay plate. The assay plates were prepared by first preparing YPAD containing 1% glucose and the highest concentration of each media type (fluconazole = 4 µg/mL; micafungin = 0.128 µg/mL; SDS = 0.24%) and then performing five 2-fold serial dilutions.

For broth microdilution assays, inoculated assay plates were incubated at 30°C in a humidified chamber without shaking. Cells were resuspended by pipetting and OD_600_ measured in an BioTek Epoch 2 microplate reader. Bar plots show the OD_600_ relative to the OD_600_ in YPAD (1% dextrose) without drug at or 2-fold below the parental strain MIC_50_, defined as the drug concentration that results in at least 50% reduction in growth relative to rich media. All assays were performed in triplicate.

For growth assays, inoculated assay plates with sterilized water motes surrounding the outer wells (Thermo Scientific NUNC Edge, Cat. No 267427) were grown in a BioTek Epoch 2 at 30°C with shaking for 48 hours, with OD_600_ readings taken every 15 minutes.

### Spot plates

Cells were patched from glycerol stocks onto solid YPAD agar and grown at 30°C for 1 day. Liquid YPAD was then inoculated with a small amount of cells from each patch and grown overnight at 30°C with shaking. Cells were serially diluted to 10^5^-10^2^ cells/mL in PBS and 5 µL of each dilution was spotted onto to YPAD agar plates containing 1% glucose and the concentration of either calcofluor white or Congo red described in each figure. Plates were incubated at 30°C for 24 hours for calcofluor white and 48 hours for Congo red and imaged using a BioRad Molecular Imager Gel Doc XR+ with a black background and epi-white illumination.

### Growth curve assays for CNV strains and progenitors

Previously evolved CNV strains, isolated single colonies, and their euploid progenitors^20,37,38^ were grown up from glycerol stock in YPAD. After overnight growth, all wells were diluted 10x and OD_600_ was measured using an Epoch 2 plate reader. Each well was then normalized to an OD of 0.01 and used to inoculate three plates, all of which contained YPAD with 1% glucose, resulting in a final starting OD of 0.001. Two plates contained YPAD alone and each of the environmental condition additive in each row, and one plate contained only YPAD with 1% glucose that would be grown at 37°C. Each plate was placed in a BioTek Epoch 2 and grown at 30°C or 37°C with shaking for 48 hours, with OD_600_ measurements taken every 15 minutes. Data was processed in R (v 4.4.0). Area under the curve was calculated from empirical measurements using the package growthcurver^64^ and each environmental condition was first made relative to the rich medium condition (YPAD) by subtracting growth in YPAD from growth in the stress condition, and then differences between CNV strains and their matched progenitor strain again calculated by subtracting the progenitor from the CNV strain for each replicate.

### RT-qPCR

Individually generated CRISPRa transformants and their progenitors were patched onto solid YPAD agar from glycerol stocks and grown for 2 days at 30°C. 3 mL of liquid YPAD was inoculated with a small portion of the patch and grown overnight at 30°C with shaking. 350 µL of the overnight culture was used to inoculate 50 mL of YPAD. When the cultures reached an OD between 0.4-0.5, each culture was pelleted, supernatant removed, and cell pellets flash frozen in liquid nitrogen. Cell pellets were stored at −80°C overnight. RNA was extracted from frozen cell pellets using the Qiagen RNeasy Mini Kit (Cat No 74106), using the mechanical disruption method and an Omni BeadRuptor Elite, and on-column DNase digestion performed. cDNA was prepared using SuperScript II RT (ThermoFisher, Cat No 18064014) using Oligo(dT) primers for reverse transcripton of polyA mRNA, according to the manufacturer’s instructions. qPCR reactions were performed in technical triplicate with primers specific to each transcript (Supplementary Table S5), as well as no reverse transcriptase and no template controls, using PowerUp SYBR Green mastermix (Fisher Scientific, Cat No 425742) and run on a BioRad CFX-96. Delta-delta Ct values for each gene were calculated, with propagation of error, relative to *ACT1* (primers 680 and 681, Supplementary Table S5), and then relative to the progenitor strain.

### Predicted fitness effects of CNV strains from individual gene overexpression fitness

For each isolate containing a single CNV, the precise boundaries of the CNV were defined manually by examining whole genome sequencing read coverage across genomic regions (Supplementary Table S1). Predicted fitness effects were calculated by summing the significant fitness effects of all genes within these defined regions, regardless of region copy number, in each environmental condition relative to rich media. These sums were then compared to the growth of the actual CNV strain in the same environmental condition, after taking the difference in area under the curve of the CNV strain in the environmental condition and in rich media, and then again taking the difference between that value and the same value for the euploid progenitor strain.

### Simulated CNV fitness effects of differing sizes

Simulated CNV fitness effects were calculated by paneling across the regions of the genome targeted by CRISPRa and summing the fitness effects of all genes located within a 50Kb, 100Kb, 300Kb, or 500Kb window in fluconazole and in micafungin. Overlapping windows at 1Kb intervals were used. Code for this analysis can be found at https://github.com/selmeckilab/CRISPRactivation_for_CNV_genes.

## ACKNOWLEDGEMENTS

We thank D. Davis, W. Harcombe, C. Myers, B. Metzger and all members of the Selmecki and Myers lab for fruitful discussions on the content of this manuscript. The University of Minnesota Genomics Center and Minnesota Supercomputing Institute provided services used to conduct this work. Funding for this work was provided by the Burroughs Wellcome Fund Investigator in the Pathogenesis of Infectious Diseases Award (#1020388 to AS). Additional funding for this work was provided by the National Institutes of Health (R01AI143689 to AS and K99GM160701 to PVZ), for sequencing costs, strain engineering, strain validation, and phenotypic analyses, as well as salary support for PVZ, ES, CZ, KM, and AS. In no way were National Institutes of Health funds used to support research expenses or salary support at the University of Guelph for RSS and NCG, who were independently supported for the entirety of the project. RSS was supported by an NSERC Canada Research Chair and NCG was supported by an NSERC CGS-D award. The work of NCG and RSS was supported by CIHR Project Grant (PJT206047) and NSERC Discovery Grant (RGPIN-2026-04298) to RSS.

## SORCE CODE/DATA AVAILABILITY

All code used for data analysis and figure generation is available at https://github.com/selmeckilab/CRISPRactivation_for_CNV_genes. Raw sequencing data for all pooled bar-seq experiments and whole genome sequencing data for strains first presented in this manuscript are available at PRJNA1506084. SRA accession numbers for previously published whole genome sequencing data are listed in Supplementary Table S1. Raw barcode counts from bar-seq competition experiments are available in Supplementary Table S6, and processed gene-level fitness values for all backgrounds and environments are available in Supplementary Table S2.

## SUPPLEMENTARY FILES

**Supplementary Table S1: Strains used in this study.** List and description of all strains used in this study, including those described previously and those first generated and described in this manuscript. SRA accession numbers are included for strains that were whole genome sequenced.

**Supplementary Table S2: Fitness estimates.** A tab-delimited text file containing all final processed gene-level fitness estimates from all pooled competition assays in all environments and genetic backgrounds. The table is in long-format, and includes columns designating the overexpressed gene name in A22 format, the mean fitness values relative to rich media, the strain background, environmental condition, gene start position, chromosomal region on which the gene is located, whether the gene significantly affected fitness in at a 5% FDR in that environment, and in that genetic background, and the standard error of the mean fitness.

**Supplementary Table S3: Broth microdilution assay statistics.** A table includes the specific colony names of individually generated CRISPR-activation mutants overexpressing single genes shown in Fig.5D, the gene they target, the drug concentration shown in Fig.5D, and the p-value from the t-test comparing the growth of the CRISPR-activation strain to the growth of the progenitor strain.

**Supplementary Table S4: Pooled sgRNA sequences.** An Excel workbook with multiple sheets including gene name conversation from A21 to A22 for all genes targeted with CRISPR-activation, the list of all sgRNA sequences generated targeting each gene in the list, the addition of 3’ and 5’ overhangs for the amplification of the Twist oligo pool, and the final oligo submission for the Twist oligo pool.

**Supplementary Table S5: Primers and Plasmids used in this study.** List, sequence, and description of all primers and plasmids used in the study.

**Supplementary Table S6: All raw sgRNA counts from deep sequencing.** A data matrix of all raw barcode (sgRNA) counts from amplicon sequencing fastq files. Code generating and processing this data can be found at https://github.com/selmeckilab/CRISPRactivation_for_CNV_genes.

---

**Extended Data Fig. 1:**
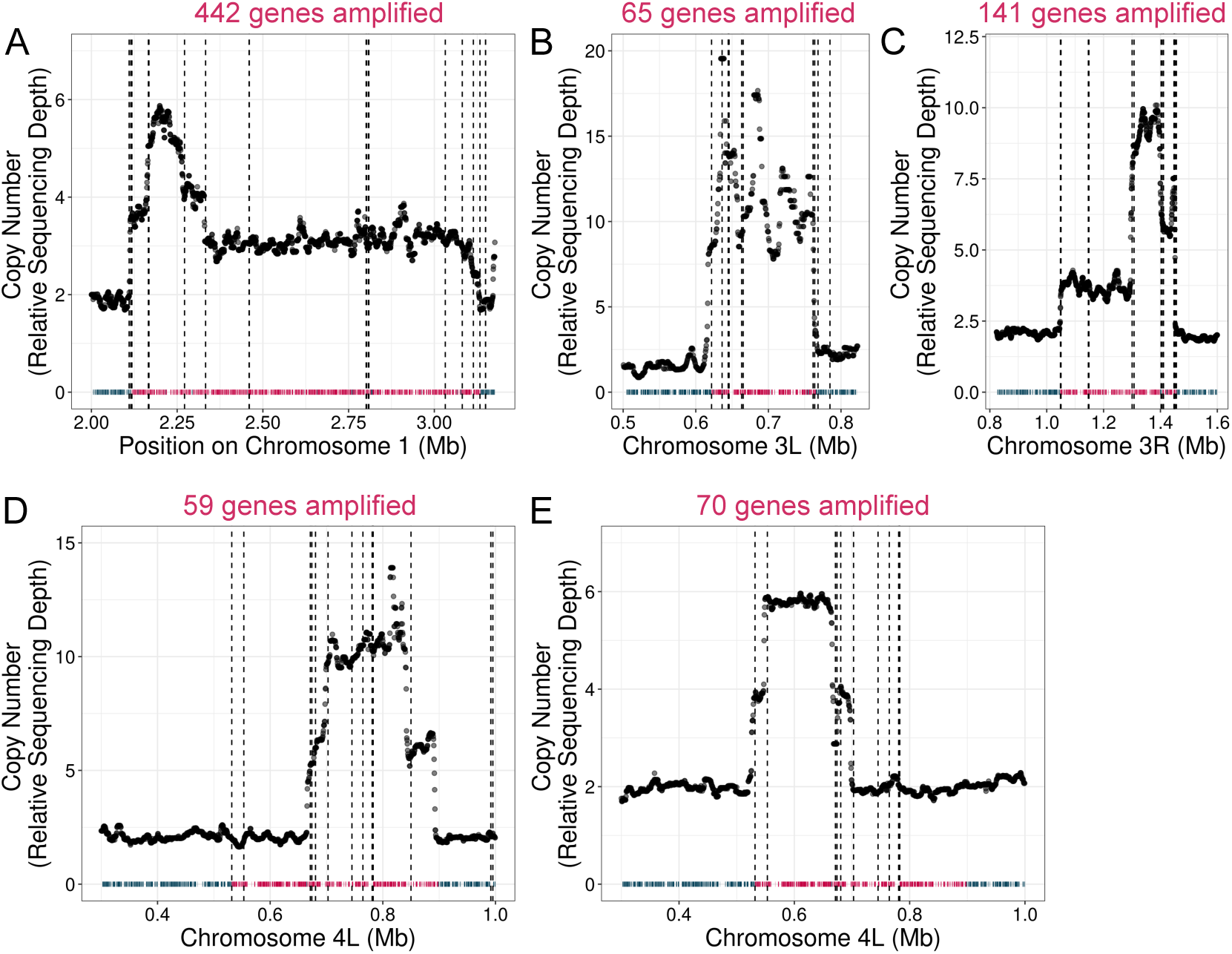
Genes in CNV regions targeted for overexpression by CRISPR-activation. Whole genome sequencing read depth of five azole-evolved strains with segmental chromosome amplifications. Read depth is plotted on the y-axis as a rolling median across chromosome position. Each panel depicts whole genome sequencing data for a different evolved isolate with CNVs on (A) chromosome 1 (AMS4105), (B) the left arm of chromosome 3 (AMS3092), (C) the right arm of chromosome 3 (AMS4105), (D, E) the left arm of chromosome 4 (AMS4702 and AMS5778, respectively). Each plot also shows the position of each encoded gene as a tick mark along the x-axis. The number of genes amplified in the CNV is written above each plot. Genes in red are those targeted by the CRISPR-activation system in the euploid CRISPRa pools. Dotted lines are positions of repeat elements with a high sequence identity partner in the CNV region, as defined in Todd et al, 2019. The two CNVs on chromosome 4 are partially overlapping, and all genes in both CNVs are targeted, resulting in a larger area targeted than covered by each individual CNV. More strain information can be found in Supplementary Table S1.

**Extended Data Fig. 2:**
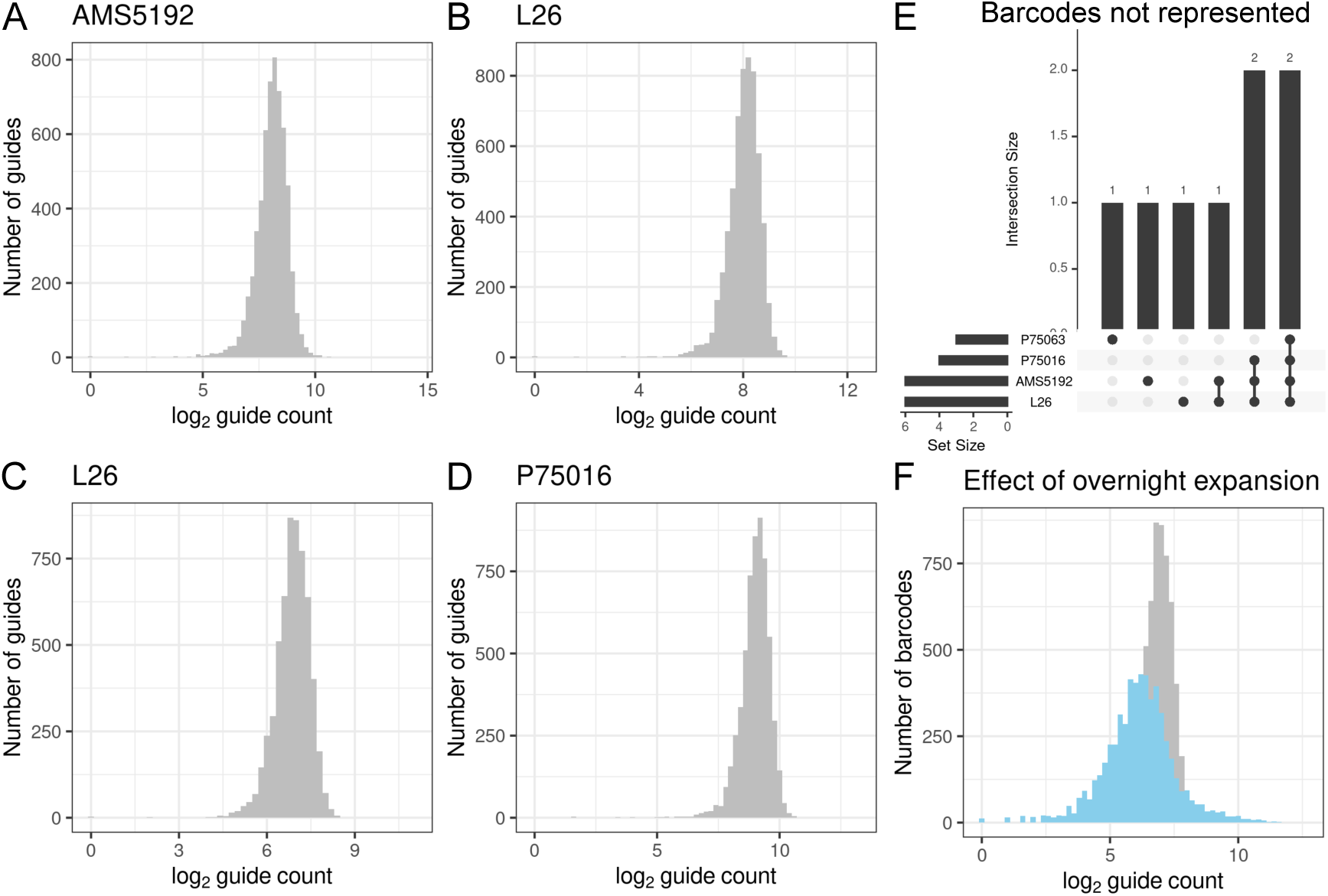
Barcode composition of the four fungal pools. Histograms of log_2_ guide counts for the fungal pools generated in four different genetic backgrounds: (A) AMS5192, (B) L26, (C) P75063, and (D) P75016. Each pool shows a similarly shaped distribution of guide frequencies. (E) Comparison of guides that are not detected in each pool. Total number of undetected guides is shown by the bar chart on the left of each pool name. Overlap between each pool is indicated by the barchart above the dots connecting the pools in which that set of guides is not detected. (F) A histogram of guide frequencies detected in the P75063 pool before (grey) and after (blue) an overnight expansion in rich media. The change in barcode frequencies in this constitutively expressed system during an overnight expansion led us to perform a short recovery (5hrs) immediately before competition rather than a full overnight culture.

**Extended Data Fig. 3:**
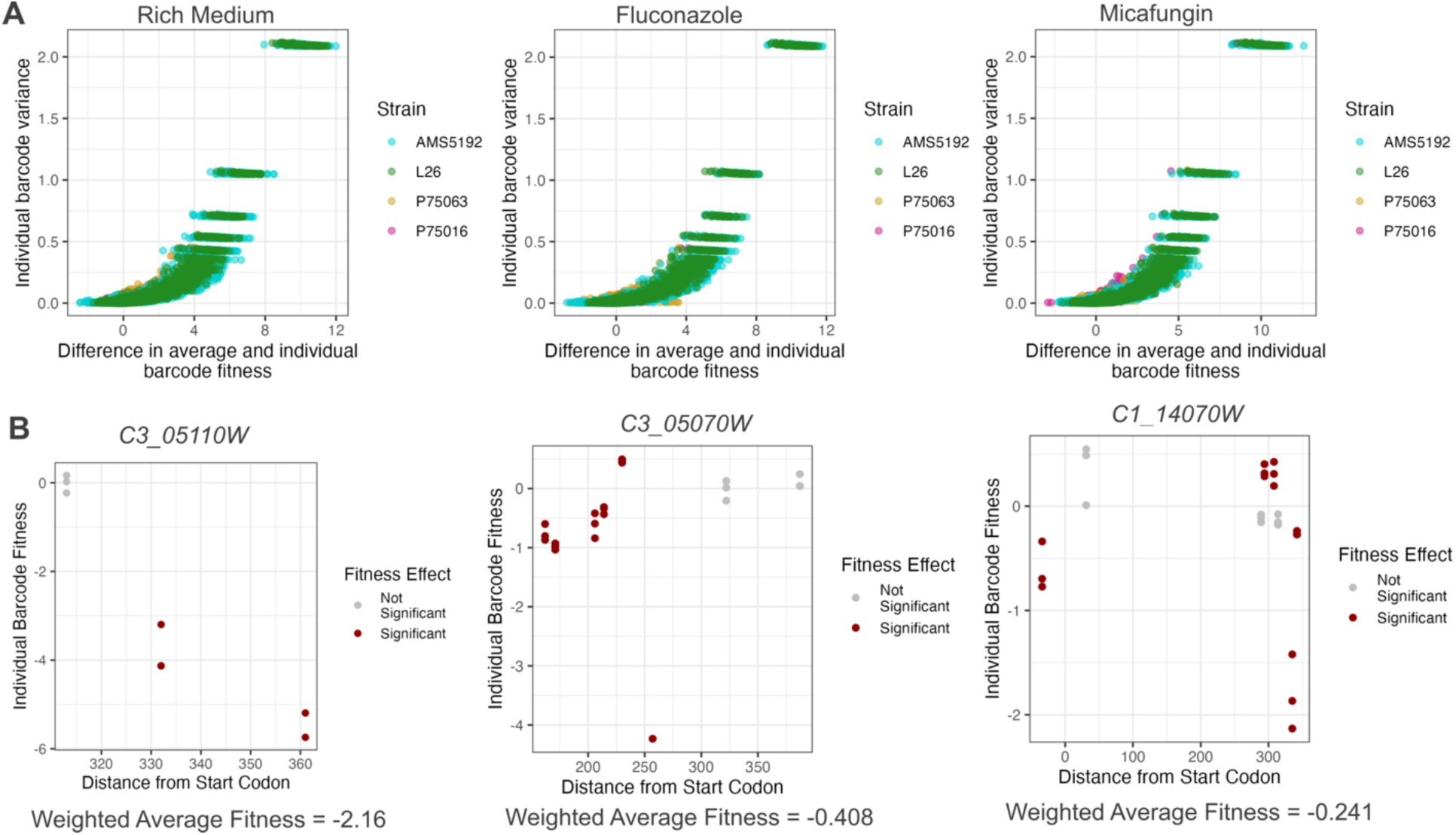
Gene-level weighted average fitness values are more robust and conservative than individual sgRNA fitness values. (A) For each individual sgRNA strain, the absolute difference between the individual sgRNA fitness score and the weighted average fitness score for all sgRNAs targeting the same gene is plotted on the x-axis and the variance for the individual barcoded strain is plotted on the y-axis. All barcoded strains are shown for competition in rich medium (left), fluconazole (center), and micafungin (right) and colored according to each genetic background. (B) Each panel shows the fitness estimates for each sgRNA targeting a single gene in rich medium plotted according to their distance from the start codon of the gene they target on the y-axis. Each sgRNA strain is shown three times, once for each experimental replicate. Individual sgRNA strains that have significantly different fitness from the non-targeting controls in rich medium at a 5% FDR are shown in red. On the left is *C3_05110W*, with three guides that have relatively consistent negative fitness effects and result in a weighted average fitness effect of −2.16. In the center is *C3_05070W*, which has more variable fitness estimates between sgRNAs, including one barcoded strain with significant positive fitness effects and one outlying barcoded strain with strong negative fitness effects. Multiple barcoded strains with smaller negative fitness effects results in a weighted average fitness effect of −0.408. On the right is *C1_14070W*, which has variable effects between barcoded strains targeting regions very close to each other and results in a weighted average fitness of −0.241. See Methods section “Calculation of gene-level fitness values” for additional detail.

**Extended Data Fig. 4:**
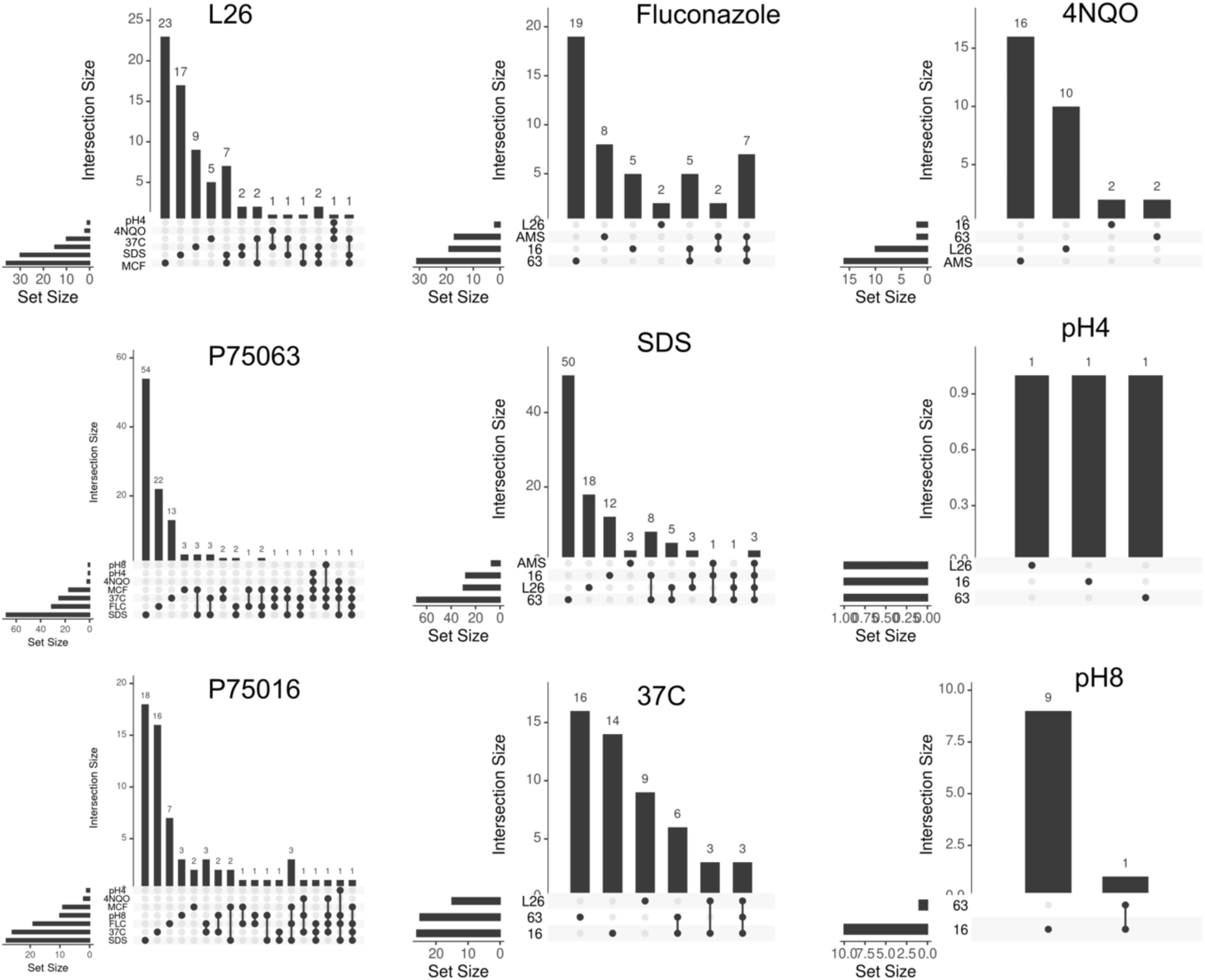
Upset plots showing overlapping sets of genes with significant fitness effects across genetic backgrounds and environments. Each plot shows the total number of genes with significant fitness effects in the environmental condition or genetic background indicated by the horizontal barchart to the left of each condition name. Genetic background names have been abbreviated (AMS5192 = AMS, P75063 = 63, P75016 = 16, and L26 = L26). Set intersections are shown as vertical columns on the right of each plot, and the identity of each intersection indicated by a black point below. Genes that significantly affect fitness in only condition are indicated with a single dot, while intersections of multiple conditions are indicated by dots connected with a black line.

**Extended Data Fig. 5:**
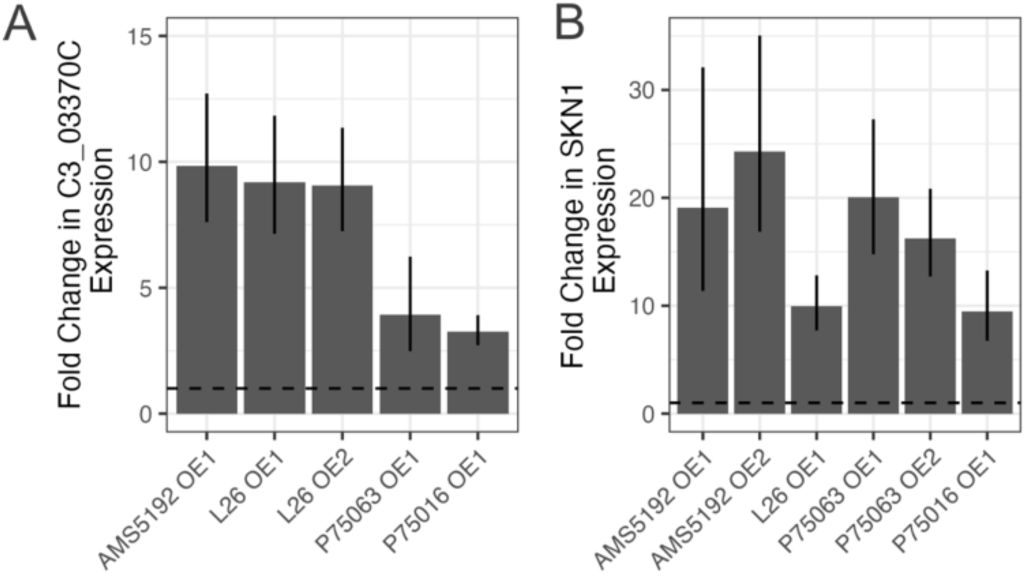
RT-qPCR of select independently generated CRISPRa transformants. (A) Expression change in *C3_03370C* as measured via RT-qPCR relative to each wild type progenitor in all four genetic backgrounds when transformed to contain the CRISPRa construct with the same sgRNA targeting *C3_03370C*. All samples are first normalized to *ACT1* expression. Multiple independent mutants containing the same sgRNA are shown for the L26 background. A dotted line showing a log fold change of 1 (no change relative to the untransformed progenitor) is represented with a dotted line. Bars are the mean of three replicates and whiskers are standard deviations with propagation of error. (B) As in (A), but for expression of *SKN1* in transformants with sgRNA targeting *SKN1*. Multiple independent transformants are shown for AMS5192 and P75063.

**Extended Data Fig. 6:**
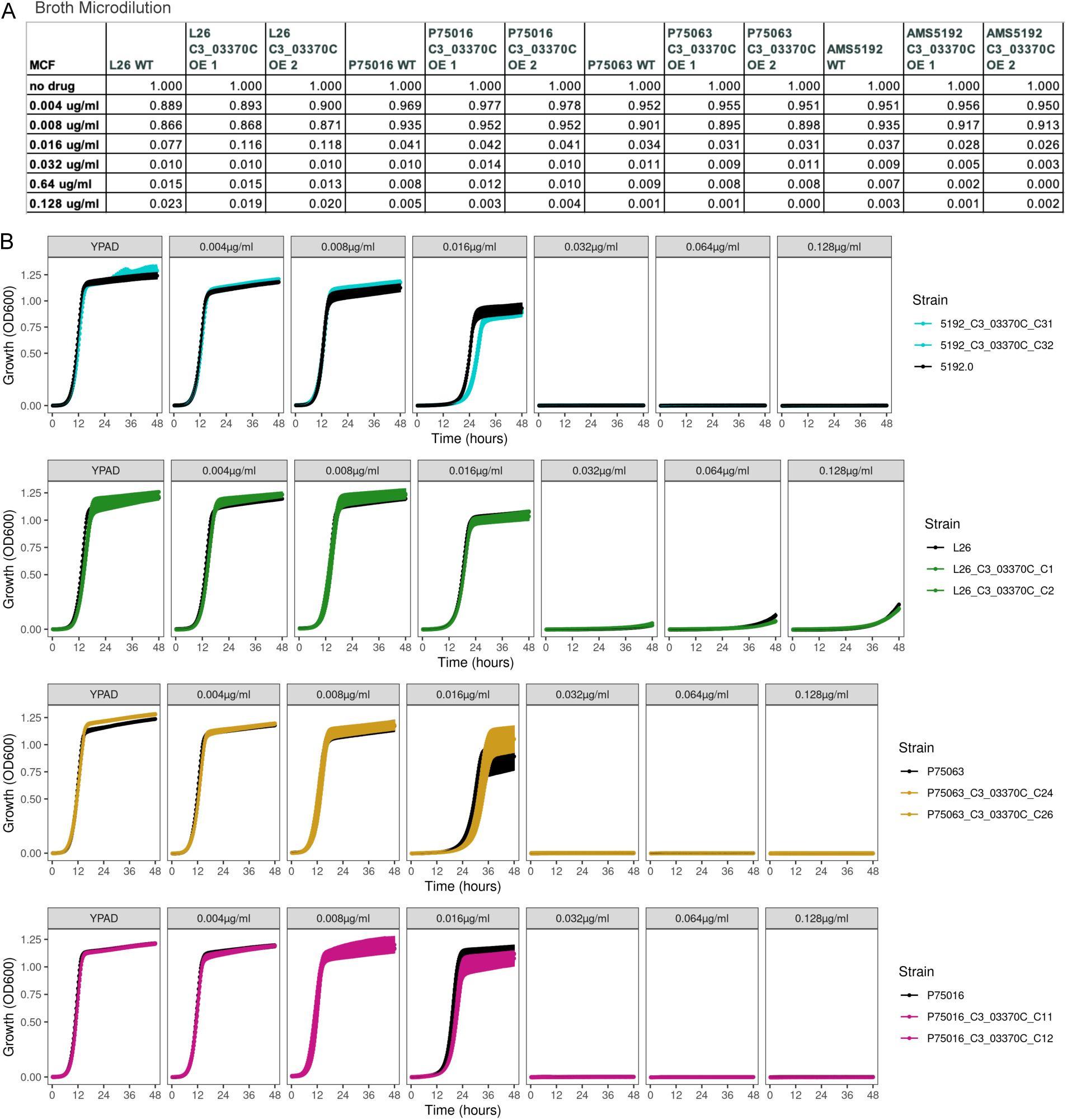
***C3_03370C* overexpression phenotypes in micafungin.** (A) From a static microbroth dilution assay, a table indicates the relative OD_600_ values of each strain shown in the column in the drug concentration listed in each row relative to 0 µg/mL micafungin. Two independent CRISPRa mutants were assayed for each genetic background. (B) Shaking growth curve assays show OD_600_ measured in 15 minute intervals in rich media (YPAD) or at varying concentrations of micafungin for the wild type genetic background (black) and two independently generated CRISPRa transformants overexpressing *C3_03370C,* colored according to the genetic background. Independent transformants are named as described in Supplementary Table S1.

**Extended Data Fig. 7:**
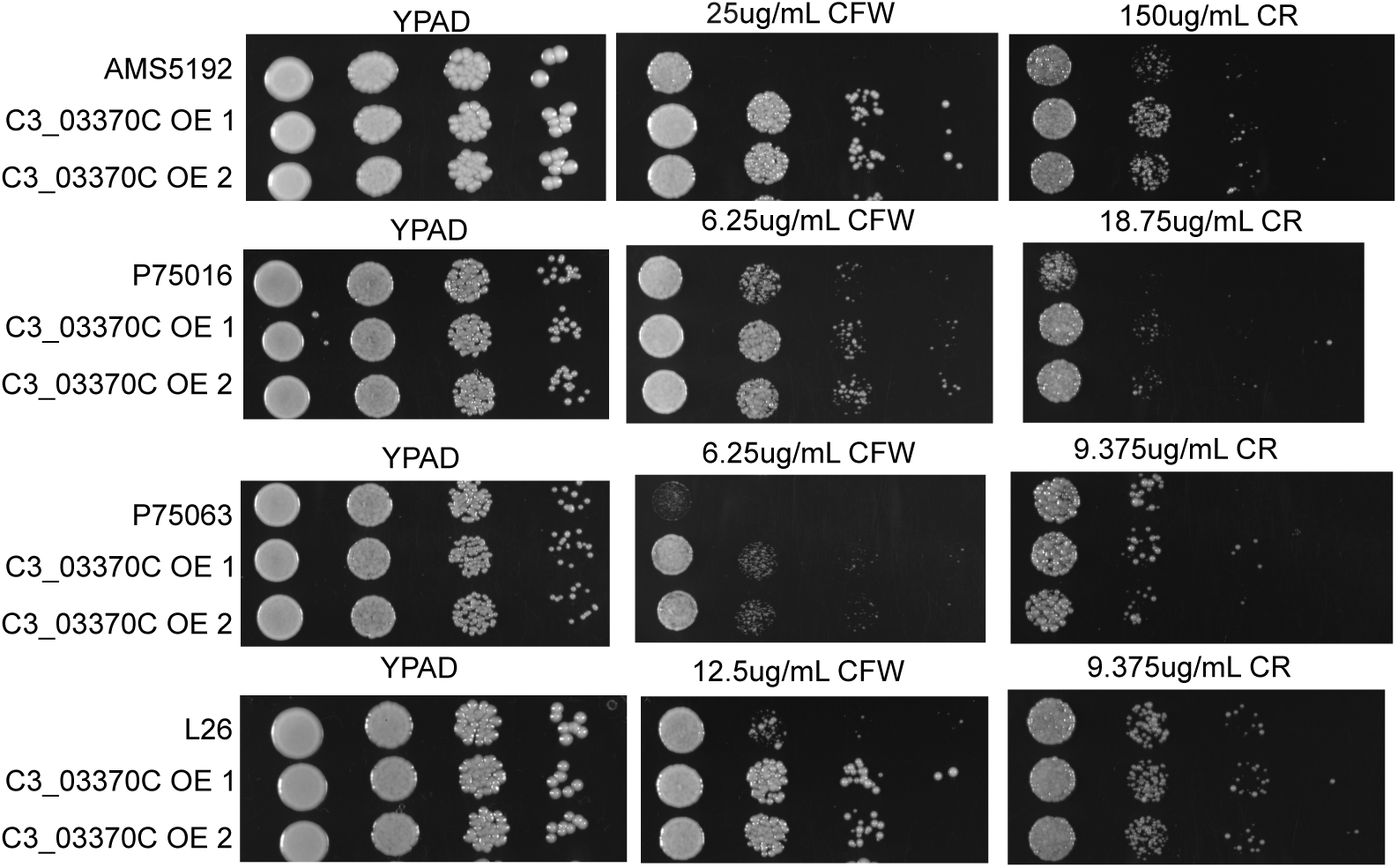
Spot Plates. As in Figure 4D, cells spotted onto agar containing either YPAD media with no additives, calcofluor white (CFW), or Congo red (CR) from left to right are shown after 24 (CFW) or 48 (CR) hours of growth. Spots are at cell densities of 10^5^, 10^4^, 10^3^, and 10^2^ cells/mL from left to right. Progenitor strains (top rows) are compared to two independently generated CRISPRa mutants overexpressing *C3_03370C* (middle and bottom rows). Spots from AMS5192 and P75063 are repeated from Figure 4D for comparison to the additional two backgrounds.

**Extended Data Fig. 8:**
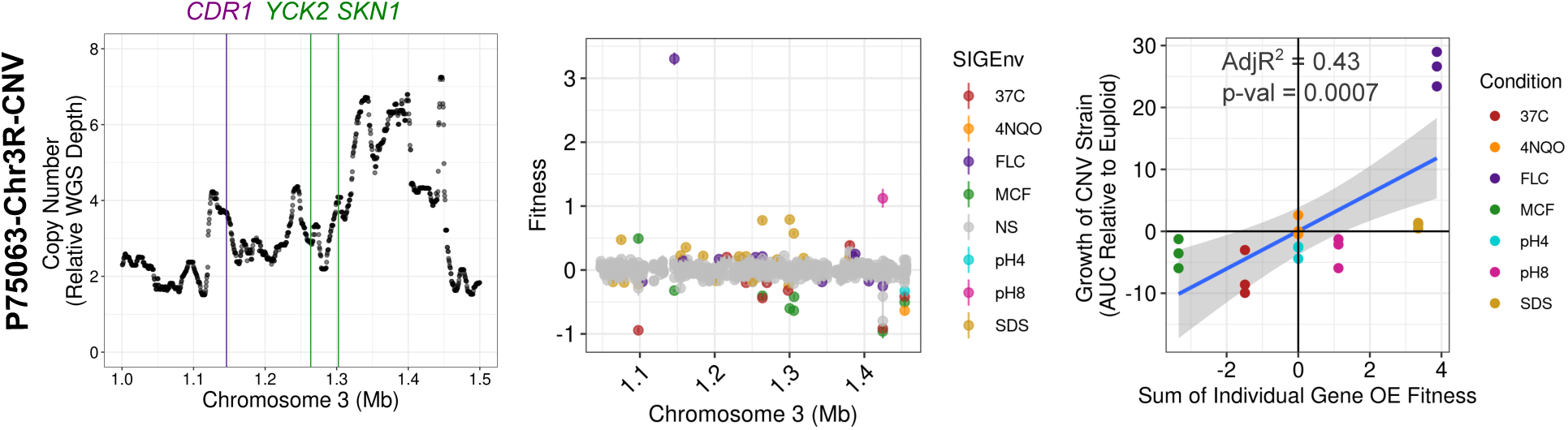
Single-gene fitness predictions for P75063-Chr3R-CNV. As in figure 6, a rolling median of mean whole genome sequencing read depth normalized to median read depth is plotted on the y-axis across the genomic positions shown on the x-axis for a region of chromosome 3R in strain P75063-Chr3R-CNV. Vertical lines are the positions of the genes *CDR1*, *YCK2*, and *SKN1*. (B) Fitness of single gene overexpression from the CRISPRa bulk competition assay is shown for the region of chromosome 3R amplified in strain P75063-Chr3R-CNV. Each gene is shown multiple times for each environmental condition, but only background P75063 is shown. Genes without significant fitness effects are shown in grey, while genes with significant fitness effects are colored according to the environment in which they have the effect. Points are means of three replicates and bars are standard error of the mean. (C) Relative growth, as measured by the area under the curve (AUC) of strain P75063-Chr3R-CNV relative to its matched progenitor strain P75063, is plotted on the y-axis. Relative growth in each condition relative to rich media is shown in different colors as indicated in the legend. Triplicate measures of CNV growth are shown as individual points. Predicted fitness effect by summing the significant fitness effects of genes in the amplified region are plotted on the x-axis. Because there is only one predicted fitness by sum, but replicate measures for the CNV strain, there is one x-axis value for each of three y-axis measures.

**Extended Data Fig. 9:**
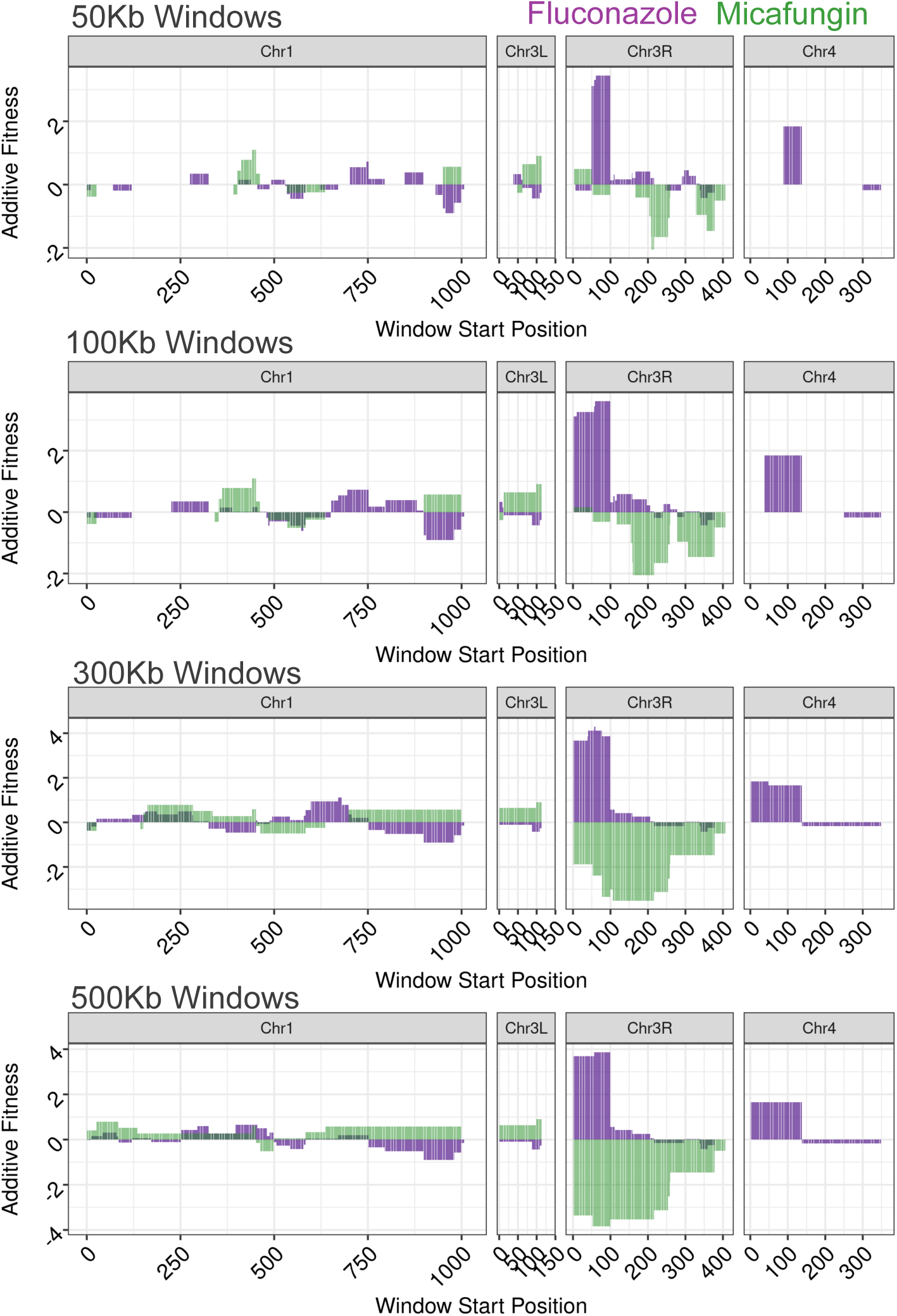
Simulated CNV fitness in fluconazole and micafungin for all CNV regions and window sizes. Predicted fitness effect of simulated CNVs of 50Kb, 100Kb, 300Kb, and 500Kb, paneling at 1Kb intervals across the regions of chromosome 1, chromosome 3L, chromosome 3R, or chromosome 4 targeted by CRISPRa. Fitness effects were predicted by adding individual gene fitness in each region in fluconazole (purple) or micafungin (green) in background P75063.

**Extended Data Fig. 10:**
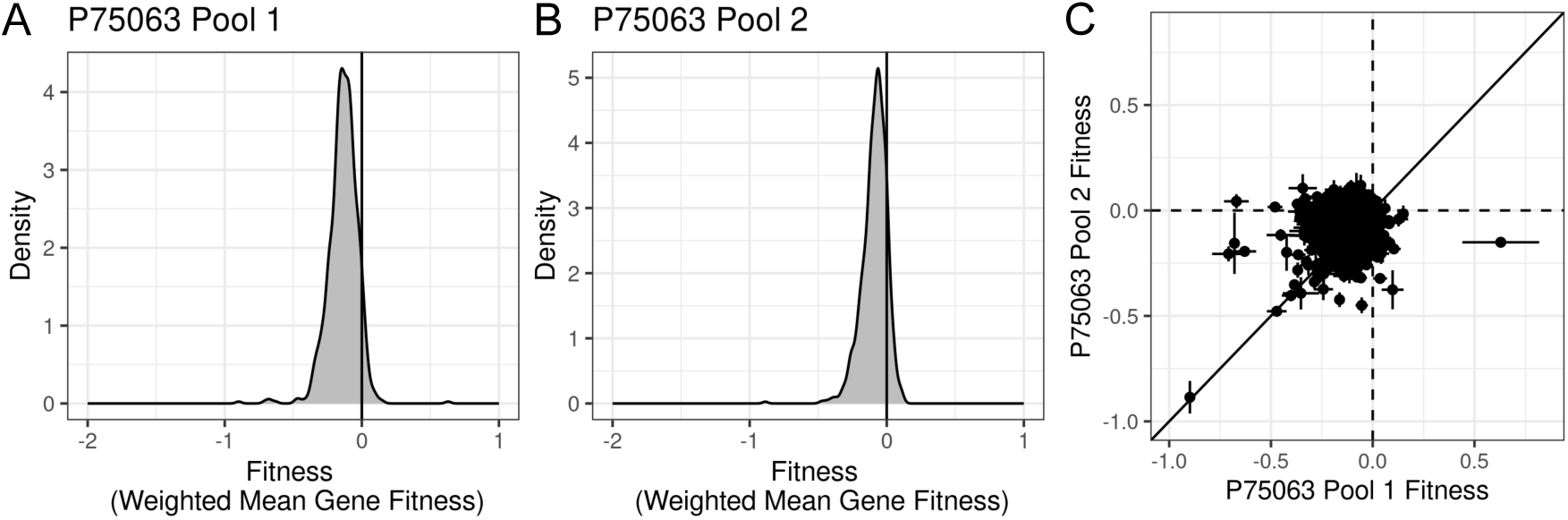
Individually generated pool comparison for P75063. (A) A density plot of the distribution of fitness effects of overexpression for all genes in rich media for the first pool generated in the genetic background P75063. (B) A density plot showing the distribution of fitness effects of overexpression for all genes in rich media for the second, independently generated fungal pool in the genetic background P75063. (C) A scatterplot shows the fitness effects of overexpression in rich media for each gene (points) in the first pool generated in P75063 (x-axis) and the second pool generated in P75063 (y-axis). X = y line is in black, x- and y-intercepts are shown as dotted lines, points are means of three replicates, and error bars are standard error of the mean.

## REFERENCES

1. Fritts, R. K., Ebmeier, C. C. & Copley, S. D. Transcriptomic and proteomic ramifications of segmental amplification in Escherichia coli. Proc. Natl. Acad. Sci. U. S. A. 122, e2422424122 (2025).

2. Nair, S. et al. Adaptive copy number evolution in malaria parasites. PLoS Genet. 4, e1000243 (2008).

3. Paulander, W., Andersson, D. I. & Maisnier-Patin, S. Amplification of the gene for isoleucyl-tRNA synthetase facilitates adaptation to the fitness cost of mupirocin resistance in Salmonella enterica. Genetics 185, 305–312 (2010).

4. Pränting, M. & Andersson, D. I. Escape from growth restriction in small colony variants of Salmonella typhimurium by gene amplification and mutation: Small colony variants of Salmonella typhimurium. Mol. Microbiol. 79, 305–315 (2011).

5. Vande Zande, P., Zhou, X. & Selmecki, A. The Dynamic Fungal Genome: Polyploidy, Aneuploidy and Copy Number Variation in Response to Stress. Annu. Rev. Microbiol. (2023) doi:10.1146/annurev-micro-041320-112443.

6. Bergin, S. et al. Analysis of clinical Candida parapsilosis isolates reveals copy number variation in key fluconazole resistance genes. Antimicrob. Agents Chemother. 68, e0161923 (2024).

7. Muenzner, J. et al. Natural proteome diversity links aneuploidy tolerance to protein turnover. Nature 630, 149–157 (2024).

8. Hose, J. et al. Dosage compensation can buffer copy-number variation in wild yeast. Elife 4, (2015).

9. Torres, E. M. et al. Effects of aneuploidy on cellular physiology and cell division in haploid yeast. Science 317, 916–924 (2007).

10. Kravets, A. et al. Widespread occurrence of dosage compensation in Candida albicans. PLoS One 5, e10856 (2010).

11. Sunshine, A. B. et al. The Fitness Consequences of Aneuploidy Are Driven by Condition-Dependent Gene Effects. PLoS Biol. 13, e1002155 (2015).

12. Yang, F. et al. The fitness costs and benefits of trisomy of each Candida albicans chromosome. Genetics 218, (2021).

13. Rojas, J. et al. Comparative modeling reveals the molecular determinants of aneuploidy fitness cost in a wild yeast model. Cell Genom. 100656 (2024).

14. Abdul-Rahman, F. & Gresham, D. Copy number variation facilitates rapid toggling between ecological strategies. bioRxivorg 2025.07.22.666191 (2025) doi:10.1101/2025.07.22.666191.

15. Lin, K. H. et al. Using antagonistic pleiotropy to design a chemotherapy-induced evolutionary trap to target drug resistance in cancer. Nat. Genet. 52, 408–417 (2020).

16. Gifford, D. R. et al. Environmental pleiotropy and demographic history direct adaptation under antibiotic selection. Heredity (Edinb.) 121, 438–448 (2018).

17. Bakerlee, C. W., Phillips, A. M., Nguyen Ba, A. N. & Desai, M. M. Dynamics and variability in the pleiotropic effects of adaptation in laboratory budding yeast populations. Elife 10, (2021).

18. Yang, F. et al. Aneuploidy Enables Cross-Adaptation to Unrelated Drugs. Mol. Biol. Evol. 36, 1768–1782 (2019).

19. Robinson, D. et al. Gene-by-environment interactions influence the fitness cost of gene copy-number variation in yeast. G3 (2023) doi:10.1093/g3journal/jkad159.

20. Todd, R. T. & Selmecki, A. Expandable and reversible copy number amplification drives rapid adaptation to antifungal drugs. Elife 9, (2020).

21. Zhou, X. et al. Single-cell detection of copy number changes reveals dynamic mechanisms of adaptation to antifungals in Candida albicans. Nat. Microbiol. 1–16 (2024).

22. Gresham, D. et al. The repertoire and dynamics of evolutionary adaptations to controlled nutrient-limited environments in yeast. PLoS Genet. 4, e1000303 (2008).

23. Robinson, D., Place, M., Hose, J., Jochem, A. & Gasch, A. P. Natural variation in the consequences of gene overexpression and its implications for evolutionary trajectories. Elife 10, (2021).

24. Selmecki, A., Forche, A. & Berman, J. Genomic plasticity of the human fungal pathogen Candida albicans. Eukaryot. Cell 9, 991–1008 (2010).

25. Hirakawa, M. P. et al. Genetic and phenotypic intra-species variation in Candida albicans. Genome Res. 25, 413–425 (2015).

26. Scott, N. E. et al. Aneuploidy, polyploidy and loss of heterozygosity distinguish serial bloodstream isolates of Candida albicans: This article is part of the Candida collection. Microb. Genom. 12, 001659 (2026).

27. Forche, A. et al. Selection of Candida albicans trisomy during oropharyngeal infection results in a commensal-like phenotype. PLoS Genet. 15, e1008137 (2019).

28. Ford, C. B. et al. The evolution of drug resistance in clinical isolates of Candida albicans. Elife 4, e00662 (2015).

29. Ropars, J. et al. Gene flow contributes to diversification of the major fungal pathogen Candida albicans. Nat. Commun. 9, 2253 (2018).

30. Selmecki, A., Forche, A. & Berman, J. Aneuploidy and isochromosome formation in drug-resistant Candida albicans. Science 313, 367–370 (2006).

31. Ene, I. V. et al. Global analysis of mutations driving microevolution of a heterozygous diploid fungal pathogen. Proc. Natl. Acad. Sci. U. S. A. 115, E8688–E8697 (2018).

32. Zheng, L., Xu, Y., Wang, C. & Guo, L. Ketoconazole induces reversible antifungal drug tolerance mediated by trisomy of chromosome R in Candida albicans. Front. Microbiol. 15, 1450557 (2024).

33. Yang, F. et al. Antifungal Tolerance and Resistance Emerge at Distinct Drug Concentrations and Rely upon Different Aneuploid Chromosomes. MBio e0022723 (2023).

34. Jay, A., Jordan, D. F., Gerstein, A. & Landry, C. R. The role of gene copy number variation in antimicrobial resistance in human fungal pathogens. npj Antimicrob Resist 3, 1–8 (2025).

35. Selmecki, A., Gerami-Nejad, M., Paulson, C., Forche, A. & Berman, J. An isochromosome confers drug resistance in vivo by amplification of two genes, ERG11 and TAC1. Mol. Microbiol. 68, 624–641 (2008).

36. Vande Zande, P., et al. Step-wise evolution of azole resistance through copy number variation followed by KSR1 loss of heterozygosity in Candida albicans. PLoS Pathog. 20, e1012497 (2024).

37. Todd, R. T. et al. Antifungal drug concentration impacts the spectrum of adaptive mutations in Candida albicans. Mol. Biol. Evol. (2023) doi:10.1093/molbev/msad009.

38. Todd, R. T., Wikoff, T. D., Forche, A. & Selmecki, A. Genome plasticity in Candida albicans is driven by long repeat sequences. Elife 8, (2019).

39. Gervais, N. C. et al. Development and applications of a CRISPR activation system for facile genetic overexpression in Candida albicans. G3 13, (2023).

40. Gervais, N. C. et al. Chromosome-scale CRISPR screening reveals secretory pathway genes as drivers of aneuploidy-mediated antifungal tolerance. bioRxiv 2026.07.23.740334 (2026) doi:10.64898/2026.07.23.740334.

41. Wu, W., Lockhart, S. R., Pujol, C., Srikantha, T. & Soll, D. R. Heterozygosity of genes on the sex chromosome regulates Candida albicans virulence. Mol. Microbiol. 64, 1587–1604 (2007).

42. Wang, J. M. et al. Intraspecies Transcriptional Profiling Reveals Key Regulators of Candida albicans Pathogenic Traits. MBio 12, (2021).

43. Cravener, M. V. et al. Reinforcement amid genetic diversity in the Candida albicans biofilm regulatory network. PLoS Pathog. 19, e1011109 (2023).

44. Gerstein, A. C. & Berman, J. Candida albicans Genetic Background Influences Mean and Heterogeneity of Drug Responses and Genome Stability during Evolution in Fluconazole. mSphere 5, (2020).

45. Prasad, R., Rawal, M. K. & Shah, A. H. Candida Efflux ATPases and Antiporters in Clinical Drug Resistance. Adv. Exp. Med. Biol. 892, 351–376 (2016).

46. Aggeli, D., Li, Y. & Sherlock, G. Changes in the distribution of fitness effects and adaptive mutational spectra following a single first step towards adaptation. Nat. Commun. 12, 5193 (2021).

47. Kryazhimskiy, S., Rice, D. P., Jerison, E. R. & Desai, M. M. Global epistasis makes adaptation predictable despite sequence-level stochasticity. Science 344, (2014).

48. Askew, C. et al. Transcriptional regulation of carbohydrate metabolism in the human pathogen Candida albicans. PLoS Pathog. 5, e1000612 (2009).

49. Carolus, H. et al. Collateral sensitivity counteracts the evolution of antifungal drug resistance in Candida auris. Nat. Microbiol. 9, 2954–2969 (2024).

50. Wensing, L. F. et al. Pooled CRISPRi screening reveals fungal-specific drug target candidates. Nat. Microbiol. (2026) doi:10.1038/s41564-026-02356-w.

51. Holmes, A. R. et al. Heterozygosity and functional allelic variation in the Candida albicans efflux pump genes CDR1 and CDR2. Mol. Microbiol. 62, 170–186 (2006).

52. Coste, A. et al. A Mutation in Tac1p, a Transcription Factor Regulating CDR1 and CDR2, Is Coupled With Loss of Heterozygosity at Chromosome 5 to Mediate Antifungal Resistance in Candida albicans. Genetics 172, 2139–2156 (2006).

53. Cowen, L. E., Sanglard, D., Howard, S. J., Rogers, P. D. & Perlin, D. S. Mechanisms of Antifungal Drug Resistance. Cold Spring Harb. Perspect. Med. 5, a019752–a019752 (2014).

54. Kohanovski, I. et al. Aneuploidy Can Be an Evolutionary Diversion on the Path to Adaptation. Mol. Biol. Evol. 41, (2024).

55. Wensing, L., et al. Pooled CRISPRi and CRISPRa strain library build for Candida albicans. protocols.io https://www.protocols.io/view/pooled-crispri-and-crispra-strain-library-build-fo-6qpvrb16plmk/v1 (2026).

56. Skrzypek, M. S. et al. The Candida Genome Database (CGD): incorporation of Assembly 22, systematic identifiers and visualization of high throughput sequencing data. Nucleic Acids Res. 45, D592–D596 (2017).

57. Zhang, J., Kobert, K., Flouri, T. & Stamatakis, A. PEAR: a fast and accurate Illumina Paired-End reAd mergeR. Bioinformatics 30, 614–620 (2014).

58. Tange, O. GNU Parallel 20210822 (’Kabul’). (Zenodo, 2021). doi:10.5281/ZENODO.5233953.

59. Martinson, J. N. V. et al. Mutualism reduces the severity of gene disruptions in predictable ways across microbial communities. ISME J. 17, 2270–2278 (2023).

60. Bolger, A. M., Lohse, M. & Usadel, B. Trimmomatic: a flexible trimmer for Illumina sequence data. Bioinformatics 30, 2114–2120 (2014).

61. Li, H. Aligning sequence reads, clone sequences and assembly contigs with BWA-MEM. arXiv [q-bio.GN] (2013).

62. Li, H. et al. The Sequence Alignment/Map format and SAMtools. Bioinformatics 25, 2078– 2079 (2009).

63. Ushey, K. RcppRoll: Efficient Rolling / Windowed Operations. Preprint at https://CRAN.R-project.org/package=RcppRoll (2018).

64. Sprouffske, K. & Wagner, A. Growthcurver: an R package for obtaining interpretable metrics from microbial growth curves. BMC Bioinformatics 17, 172 (2016).

